# Inactive Parp2 causes Tp53-dependent lethal anemia by blocking replication-associated nick ligation in erythroblasts

**DOI:** 10.1101/2024.03.12.584665

**Authors:** Xiaohui Lin, Dipika Gupta, Alina Vaitsiankova, Seema Khattri Bhandari, Kay Sze Karina Leung, Demis Menolfi, Brian J. Lee, Helen R. Russell, Steven Gershik, Wei Gu, Peter J. McKinnon, Françoise Dantzer, Eli Rothenberg, Alan E. Tomkinson, Shan Zha

## Abstract

PARP1&2 enzymatic inhibitors (PARPi) are promising cancer treatments. But recently, their use has been hindered by unexplained severe anemia and treatment-related leukemia. In addition to enzymatic inhibition, PARPi also trap PARP1&2 at DNA lesions. Here, we report that unlike *Parp2^-/-^* mice, which develop normally, mice expressing catalytically-inactive Parp2 (E534A, *Parp2^EA/EA^*) succumb to *Tp53-* and *Chk2*-dependent erythropoietic failure *in utero*, mirroring *Lig1^-/-^* mice. While DNA damage mainly activates PARP1, we demonstrate that DNA replication activates PARP2 robustly. PARP2 is selectively recruited and activated by 5’-phosphorylated nicks (5’p-nicks) between Okazaki fragments, typically resolved by Lig1. Inactive PARP2, but not its active form or absence, impedes Lig1- and Lig3-mediated ligation, causing dose-dependent replication fork collapse, particularly harmful to erythroblasts with ultra-fast forks. This PARylation-dependent structural function of PARP2 at 5’p-nicks explains the detrimental effects of PARP2 inhibition on erythropoiesis, revealing the mechanism behind the PARPi-induced anemia and leukemia, especially those with TP53/CHK2 loss.

**Significance:** This work shows that the hematological toxicities associated with PARP inhibitors stem not from impaired PARP1 or PARP2 enzymatic activity but rather from the presence of inactive PARP2 protein. Mechanistically, these toxicities reflect a unique role of PARP2 at 5’-phosphorylated DNA nicks during DNA replication in erythroblasts.

## Introduction

Poly ADP-ribose polymerase (PARP) 1 and 2 are sensors of DNA strand breaks. DNA damage transiently recruits and allosterically activates PARP1 and 2. Active PARP1 and PARP2 use NAD+ as the substrate to generate poly(ADP-ribose) (PAR) chains on themselves and other proteins (*e.g.*, histone)^1^. The PAR chains, which bear a highly negative charge, promote chromatin relaxation and recruit other DNA repair proteins, including the XRCC1-DNA Ligase 3 (LIG3) complex^2^ implicated in single-strand DNA (ssDNA) break repair. In 2005, two groups reported that BRCA1/2-deficient cancer cells are extremely hypersensitive to NAD+ competitive inhibitors of PARP1 and 2^3,4^. Since then, the FDA has approved four PARP inhibitors (PARPi) for treating BRCA-deficient ovarian, breast, prostate, and pancreatic cancers. However, in 2022, the FDA withdrew two of these four PARPi from maintenance therapies owing to severe anemia and a significant increase in treatment-related myeloid leukemia^5^. Analyses of cancer patients before and after treatment also showed that PARPi induces stronger clonal hematopoiesis (CH) than canonical genotoxic therapies (*e.g.*, cisplatin)^6^. CH, especially those with Tp53 deficiency, has been linked with an increased risk for myeloid leukemia^7^. These severe hematological toxicities were unexpected for PARPi given that complete loss of PARP1, which accounts for most (>80%) of the DNA damage-induced PARylation in cells^8^, has no impact on hematopoiesis^9,10^. *Parp2^-/-^* mice are healthy at birth and achieve normal life spans. Although older *Parp2^-/-^* mice do develop mild anemia^10,11^, suggesting a role of Parp2 in red blood cell development, it remains unclear why Parp1, as the more abundant and more active paralog, cannot compensate for the loss of Parp2 during erythropoiesis.

In addition to inhibiting PARP1/2 enzymatic activities, FDA-approved PARPi also extend the appearance of PARP1 and 2 on damaged chromatin and at laser-induced DNA damage sites – the effects that have been described colloquially as PARP “trapping”^12,13^. Consistent with “trapping,” we and others showed that FDA-approved PARPi markedly decreased the exchange of PARP2 on DNA lesions and model DNA substrates^14,15^. Curiously, however, the same PARPi do not reduce the exchange of PARP1^16,17^. Instead, the persistent PARP1 foci in the presence of PARPi reflects a continual recruitment of different PARP1 molecules to the unrepair DNA lesion due to compromised PAR-dependent recruitment of XRCC-1-Lig3 and delayed repair^16^. Correspondingly, XRCC1-deficient cells also show hyperactivation and accumulation of PARP1^16,18,19^. Nevertheless, loss of PARP1 expression confers PARPi resistance to cancer cells and patients^14,15,16,20^. In contrast to normal development of *Parp1^-/-^* mice^9,10^, mice express a catalytically inactive Parp1 heterozygously (*Parp1^+/EA^*) die *in utero* with severe genomic instability^21^. These findings, along with the moderate phenotypes of Parp1 null mice, suggest that the presence of inactive PARP proteins is critical for the therapeutic effect and, potentially, the toxicity of PARPi. In this context, loss of PARP2 protein expression alone is insufficient to compromise PARPi sensitivity in BRCA-proficient or -deficient cells^15,16^, but the physiological consequences of inactivating PARP2 remain undetermined.

Despite normal development of *Parp1^-/-^* and *Parp2^-/-^*mice, Parp1^-/-^Parp2^-/-^ mice died before E9.5^9-11^, indicating functional redundancy. However, the degree to which PARP1 and PARP2 functions overlap is not yet fully defined. Although PARP1 and PARP2 share nearly identical catalytic mechanisms, they have distinct DNA-lesion binding properties. The allosteric enzymatic activation of both proteins necessitates the direct engagement of their conserved WGR (tryptophan-glycine-arginine-rich) domain with the 5’-phosphorylated DNA ends, distinguishing DNA strand breaks over intact DNA^22-26^. In addition to the WGR domain, PARP1 also contains three Zinc Finger (ZnF) domains that collectively support nanomolar interaction with and enzymatic activation by diverse DNA lesions, including single-stranded DNA (ssDNA) nicks, gaps, and DNA double-strand breaks (DSBs) with various overhangs^22,24^. In contrast, PARP2, which lacks any ZnF domain, can only be selectively activated by 5’-phosphorylated nicks (5’p-nicks)^23,26^. Moreover, the WGR domain of PARP2 straddles the 5’p-nick and interacts with both downstream and upstream DNA^25,26^. In structural studies, two PARP2 proteins brought two blunt dsDNA together to form two 5’p-nicks with one protein bound on each nick^25,26^. This might explain why DSBs can activate PARP2 in vitro. This model also explains why expanding a 5’p-nick to 1nt-gap markedly reduces PARP2 activation^23^. The 5’p-nicks are the substrates for DNA Ligases-LIG1 and LIG3 and are the repair intermediates of both base excision repair (BER) and nucleotide excision repair (NER). Moreover, 5’p-nicks arise between all Okazaki fragments on lagging strands during normal DNA replication, where they are resolved primarily by FEN1/LIG1 mediated pathway. Consistent with the abundance of 5’p-nicks during normal replication, the PAR signal is highest in the S phase of replicating cells, and PARP1 has been proposed as a sensor of unligated Okazaki fragments that escape FEN1/LIG1 mediated repair^27,28^. Presumably, the PAR chains generated upon PARP activation recruit XRCC1-LIG3 to seal the residual nicks, constituting a backup pathway of Okazaki fragment maturation. Notably, erythropoiesis seems to be especially vulnerable to Okazaki fragment maturation defects. While Lig1 is not essential for the proliferation of other cell types tested^29^, *Lig1^-/-^* mice succumbed to lethal anemia by E16.5^30^. In this context, during the terminal differentiation phase, erythroblasts undergo a distinct replication program characterized by ultra-fast fork speeds and progressive loss of DNA methylation before enucleation^31-33^ that may especially depend on the timely processing of Okazaki fragments. Nevertheless, Parp1 null cells and mice have no major replication defects. Loss of Parp2, but not Parp1, compromises red blood cell^10^ and T cell development^34^ in adult mice. But whether and how the DNA lesion selectivity of PARP1 and PARP2 contributes to those phenotypes remains elusive.

Given the allosteric trapping of PARP2, but not PARP1, by the FDA-approved PARPi^14,15^, we investigate the physiological consequence of inactivating Parp2 by introducing a catalytically inactive mutation (E534A) into endogenous Parp2 locus. In contrast to *Parp2^-/-^* mice, *Parp2^EA/EA^* mice die shortly after E13.5 from severe anemia. Loss of *Tp53* or *Chk2*, two genes implicated in high-risk CH^7^, partially corrects the anemia and rescues the lethality, suggesting that the presence of inactive PARP2 could select for TP53/CHK2 loss in hematopoietic progenitors. Mechanistically, inactive but not active Parp2 impairs Lig1- and Lig3-mediated 5’p-nick ligation, leading to replication-associated DNA breaks, G2/M arrests, and apoptosis in fast proliferating erythroblasts. Together, these results uncovered a PARylation-dependent structure function of PARP2 in erythropoiesis that can be explained by its DNA lesion specificity, providing a mechanism for the unexpected hematological toxicities of PARPi.

## Results

### Expression of catalytically inactive Parp2 causes lethal anemia in mice

To ascertain the role of PARP2 in the clinical toxicities associated with PARP inhibition, we compared the phenotype of *Parp2^-/-^* mice to that of mice expressing an enzymatically inert Parp2 protein. To this end, we introduced the E534A mutation (corresponding to the PARylation deficient E545A mutation of human PARP2) into the endogenous mouse *Parp2* locus (*Parp2^EA^*) (fig. S1A-B). Parp2^EA^ protein is stable *in vivo* (fig. S1C) and can efficiently relocate to DNA damage sites (fig. S1D). However, consistent with its lack of PARylation activity, ectopically expressed Parp2^EA^ protein fails to recruit PAR-dependent XRCC1 to sites of DNA damage in Parp1&2 double deficient cells (fig. S1D-E). GFP-tagged Parp2^EA^ forms damage-induced foci that are markedly brighter and persist longer than those of untreated GFP-Parp2^WT^ – and, as such, resemble olaparib “trapped” GFP-Parp2^WT^ foci (fig. S1D, F, and G). Moreover, fluorescent recovery after photobleaching showed that the exchange of Parp2^EA^ on damaged DNA is also similar to olaparib-treated Parp2^WT^ and slower than that of untreated Parp2^WT^ (fig. S1H and I). These results indicated that Parp2^EA^ is allosterically trapped at DNA damage like PARPi treated Parp2^WT^.

Although *Parp2^-/-^* and *Parp2^+/EA^* mice are born at the expected ratios and develop normally (Fig. 1A and fig. S2A)^10,11^, homozygous *Parp2^EA/EA^* pups were absent at birth (Fig. 1A, p<0.0001). Instead, we found small *Parp2^EA/EA^* embryos that are pale and presumably anemic on embryonic day 13.5 (E13.5) (Fig. 1A and fig. S2B). As the primary site of E13.5 hematopoiesis, the fetal liver (FL) generates predominantly erythrocytes (TER119^+^) and some B lymphocytes (B220^+^)^31^. Strikingly, the cellularity of E13.5 FLs decreased by >70% in *Parp2^EA/EA^*mice relative to controls (Fig. 1B) due to a dramatic reduction in TER119^+^ erythrocytes (∼10-fold) and a smaller reduction in B220+ B cells (∼50%) (Fig. 1C). Thus, the proportion of erythrocytes in E13.5 FLs dropped from >70% in wildtype embryos to <40% in *Parp2^EA/EA^* embryos (Fig. 1D), resulting in a ∼3-fold decrease in the ratio of erythrocytes to B cells (fig. S2C). Although erythroid progenitors (S0, CD71^low^TER119^-^) undergoing normal mitosis were less affected, the numbers of maturing erythroblasts decreased progressively, with moderate declines in proerythroblasts (S2, CD71^+^TER119^Mid^ or stage I, CD44^High^TER119^Mid^) and greater reductions (7-10 fold) in erythroblasts (S3, CD71^+^TER119^+^ or stage II-IV, TER119^High^CD44^Low^) (Fig. 1D-E and fig. S2D). Histological analysis of *Parp2^EA/EA^* FLs revealed that the reduced levels of TER119+ cells are accompanied by high levels of TUNEL+ apoptotic cells (fig. S2E). In contrast, despite the mild anemia observed in mature *Parp2^-/-^* adults (>90 days)^10^, FL hematopoiesis in *Parp2^-/-^* embryos is largely unaffected (Fig. 1A-E and fig. S2B-E). These observations suggest that the erythropoietic defects of *Parp2^EA/EA^*FLs are caused by the presence of inactive Parp2 protein rather than the lack of Parp2 enzymatic activity.

**Figure 1.**
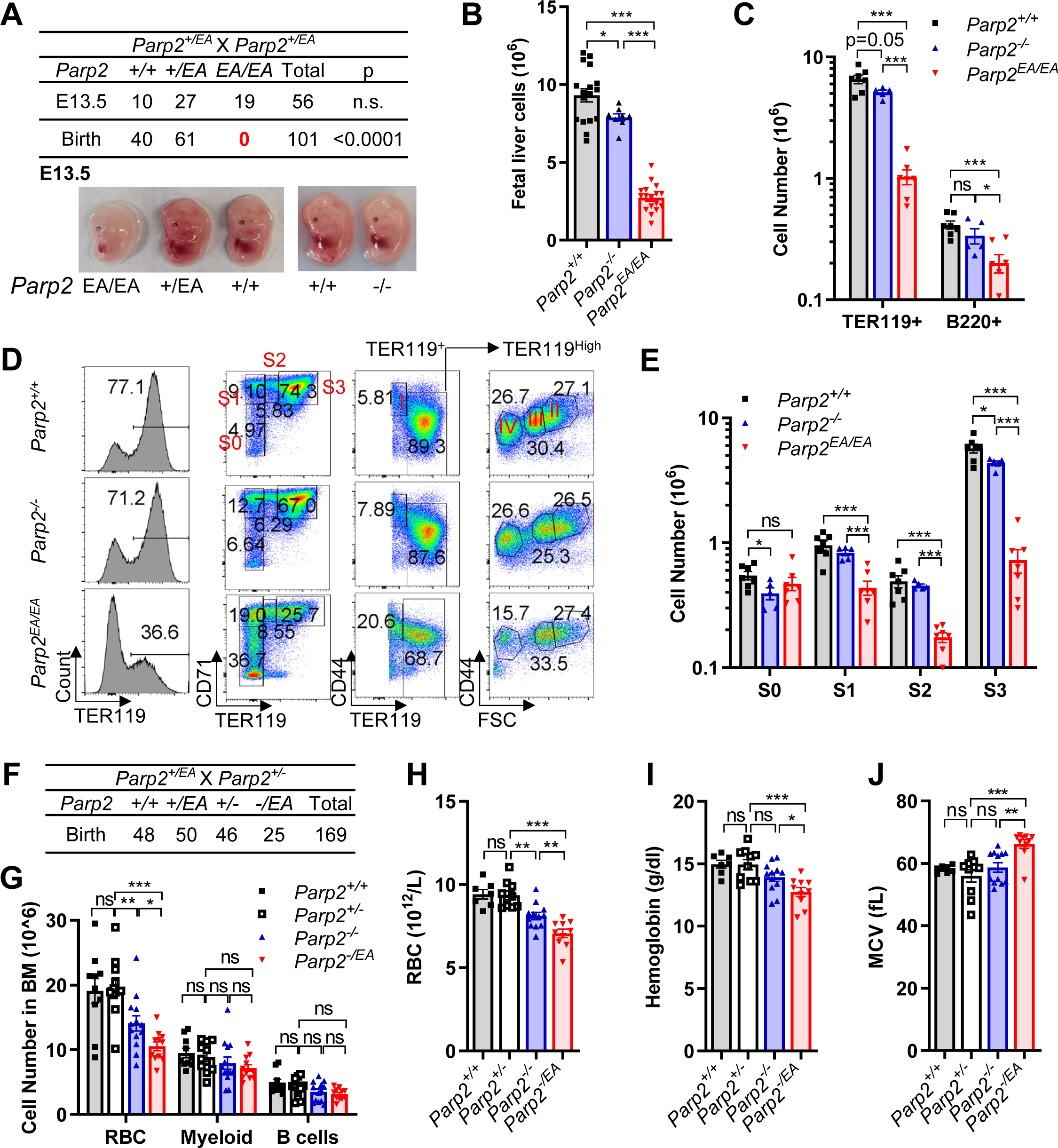
*Parp2^EA/EA^*mice are embryonic lethal with severe fetal liver erythropoiesis defects. **(A)** Upper panel: Genotype distribution from *Parp2^+/EA^*intercrossing at E13.5 and birth. Chi-squared test. n.s. p>0.05. Lower panel: Representative images of *Parp2^+/+^*, *Parp2^+/EA,^ Parp2^EA/EA^* and *Parp2^-/-^* embryos at E13.5. (**B**) Total FL cell count from E13.5 *Parp2^+/+^, Parp2^-/-^* and *Parp2^EA/EA^* embryos. (**C**) The absolute number of TER119^+^ and B220^+^ cells in the FL of E13.5 *Parp2^+/+^, Parp2^-/-^* and *Parp2^EA/EA^* embryos. (**D**) Representative flow cytometry analyses of FL cells from E13.5 *Parp2^+/+^, Parp2^-/-,^* and *Parp2^EA/EA^* embryos. The histograms of TER119 show the frequency of TER119^+^ cells of live cells. CD71 and TER119 staining determine the stage of erythropoiesis of B220^-^Thy1.2^-^Gr-1^-^CD11b^-^ live cells. S0, CD71^-^TER119^Low^; S1, CD71^+^TER119^Low^; S2, CD71^+^TER119^Mid^; S3, CD71^+^TER119^High^. CD44 and TER119 staining were followed by CD44 and FSC gating to determine the erythropoiesis differentiation stages of B220^-^Thy1.2^-^Gr-1^-^CD11b^-^TER119^+^ live cells. I, TER119^Mid^CD44^High^; (II-IV), were gated based on the progressive decrease of CD44 and FSC from TER119^High^ cells. (**E**) The absolute number of S0-S3 cells in the FL of E13.5 *Parp2^+/+^, Parp2^-/-^* and *Parp2^EA/EA^* embryos. (**F**) Genotype distribution from *Parp2^+/EA^* and *Parp2^+/-^* crossing. Chi-squared test. n.s. p>0.05. (**G**) The absolute counts of red blood cells, myeloid, and B cells in the BM from *Parp2^+/+^, Parp2^+/-,^ Parp2^-/-,^* and *Parp2^-/EA^* mice. (**H**) Red blood cell count, (**I**) White blood cell count and (**J**) Mean corpuscular volume (MCV) in peripheral blood from *Parp2^+/+^, Parp2^+/-,^ Parp2^-/-,^* and *Parp2^-/EA^* mice. The means ± SEM are shown for all the bar graphs, and an unpaired two-tailed t-test was used to measure the p values. ns, p>0.05; * p<0.05; ** p<0.01; *** p<0.001.

In support of this notion, heterozygous *Parp2^-/EA^* mice with ∼50% reduced *Parp2^EA^* protein are viable at birth (Fig. 1F and fig.S1C) but display signs of anemia that are significantly more severe than those of age-matched *Parp2^-/-^* mice, including a selective decrease in erythrocyte counts, but not lymphocyte or neutrophil counts, in both the peripheral blood and bone marrow (BM) of *Parp2^-/EA^* mice and reduced hemoglobin levels (Fig. 1G-I and fig. S3A-C). In addition, since red blood cell size decreases upon differentiation, the higher mean cell volumes (MCV) of *Parp2^-/EA^* mice (Fig. 1J) suggest the presence of immature red blood cells in the peripheral blood. Also, as erythropoietin (EPO) promotes erythropoiesis, its expression is often induced as a physiological response to anemia and anemia-associated hypoxia. Indeed, we observe elevated Epo levels in the peripheral blood of *Parp2^-/EA^* mice (fig. S3D). The anemia of *Parp2^-/EA^* mice is significantly and consistently more severe than that of age-matched *Parp2^-/-^* mice (Fig. 1G-J and fig. S3A-G). The stage-specific impairment of erythroid differentiation in the BM of *Parp2^-/EA^*adults resembles that seen in the FL of *Parp2^EA/EA^* embryos. Specifically, erythroblasts (S1-3) are reduced to a greater extent than erythrocyte progenitors (S0) in a progressive manner (fig. S3E), suggesting defects in the terminal differentiating erythroblasts. The absolute number of hematopoietic stem cells (HSC, Lin^-^Sca1^+^cKit^+^CD48^-^CD150^+^) and multi-potent progenitors (MPP, Lin^-^Sca1^+^ cKit^+^CD48^-^CD150^-^) are also slightly decreased in young adult (6-8 weeks) *Parp2^-/EA^* mice, but not *Parp2^-/-^* mice, consistent with early signs of stem cell exhaustion (fig. S3F-G). Thus, despite the difference in FL vs. BM hematopoiesis, these results indicate that the presence of inactive Parp2^EA^ protein, rather than the loss of Parp2 activity, acts in a dose-dependent manner to block erythropoiesis in both fetal liver and adult bone marrow.

### Terminally differentiating *Parp2^EA/EA^* erythroblasts display marked defects in DNA replication

How does Parp2^EA^ block erythropoiesis? Erythropoiesis can be roughly divided into two phases. In the first phase, the erythrocyte progenitors (S0) undergo largely normal mitosis to expand the numbers. In the 2^nd^ phase, the erythroblasts (from S1-S3) undergo a distinct form of rapid proliferation coupled with terminal differentiation and eventually enucleation to generate billions of new red blood cells daily. The differentiating erythroblasts are one of the most rapidly proliferating mammalian cell types with minimal interphases, evidenced by the high percentage of BrdU+ cells (>80%) in pulse labeling experiments^31,35^(Fig. 2A-B). Moreover, they also take less time in the S phase. In principle, complete replication of the same genomic DNA in a shorter duration can be achieved in two ways - firing more replication origins simultaneously (*e.g.*, Myc+ tumor cells)^36^ or increasing the speed of replication forks^31^. While most cancer cells fire additional origins driven by oncogenes and elevated CDK activities, the initiation of erythroid terminal differentiation (S0 to S1 transition) is characterized by ultra-fast progression of replication forks measured by DNA combing^31^. Consistent with those studies, both the proportions of BrdU-positive cells and the levels of BrdU label in the BrdU-positive cells are higher in S1 cells than in S0 cells from the same mice (Fig. 2A-B). In one hour, 80% of *Parp2^+/+^* and *Parp2^-/-^*FL S1-S2 cells (early erythroblasts) and >60% of S3 cells (mid-to-mature erythroblasts) are BrdU^+^ with nearly no G2/M phase (<1-2%) cells (Fig. 2A-B, p<0.001). Both BrdU^+^ cells % (S phase) and their BrdU levels (≅productive DNA synthesis per cell per hr.) dropped precipitously in *Parp2^EA/EA^* erythroblasts (Fig. 2A-C, p<0.001), indicating replication defects. Moreover, *Parp2^EA/EA^*erythroblasts display a marked activation of G1/S and G2/M checkpoints (Fig. 2D-E) and a reduced mitotic measured by histone H3S10 pho+ % (fig. S4A). In line with the relatively modest decrease of FL B cell counts (Fig. 1C), BrdU levels and cell cycle distributions are less affected in B220+ FL B cells from the same *Parp2^EA/EA^* mice (fig. S4B-D). We note that the mean BrdU levels in S1, S2, and even S3 erythroblasts are also higher than those of B220+ B cells from the same embryo (fig. S4E-F), consistent with a positive correlation between replication fork speed and vulnerability to inactive Parp2. Together, the findings suggest a model in which inactive Parp2 blocks replication in a dose-dependent and replication-dependent manner.

**Figure 2.**
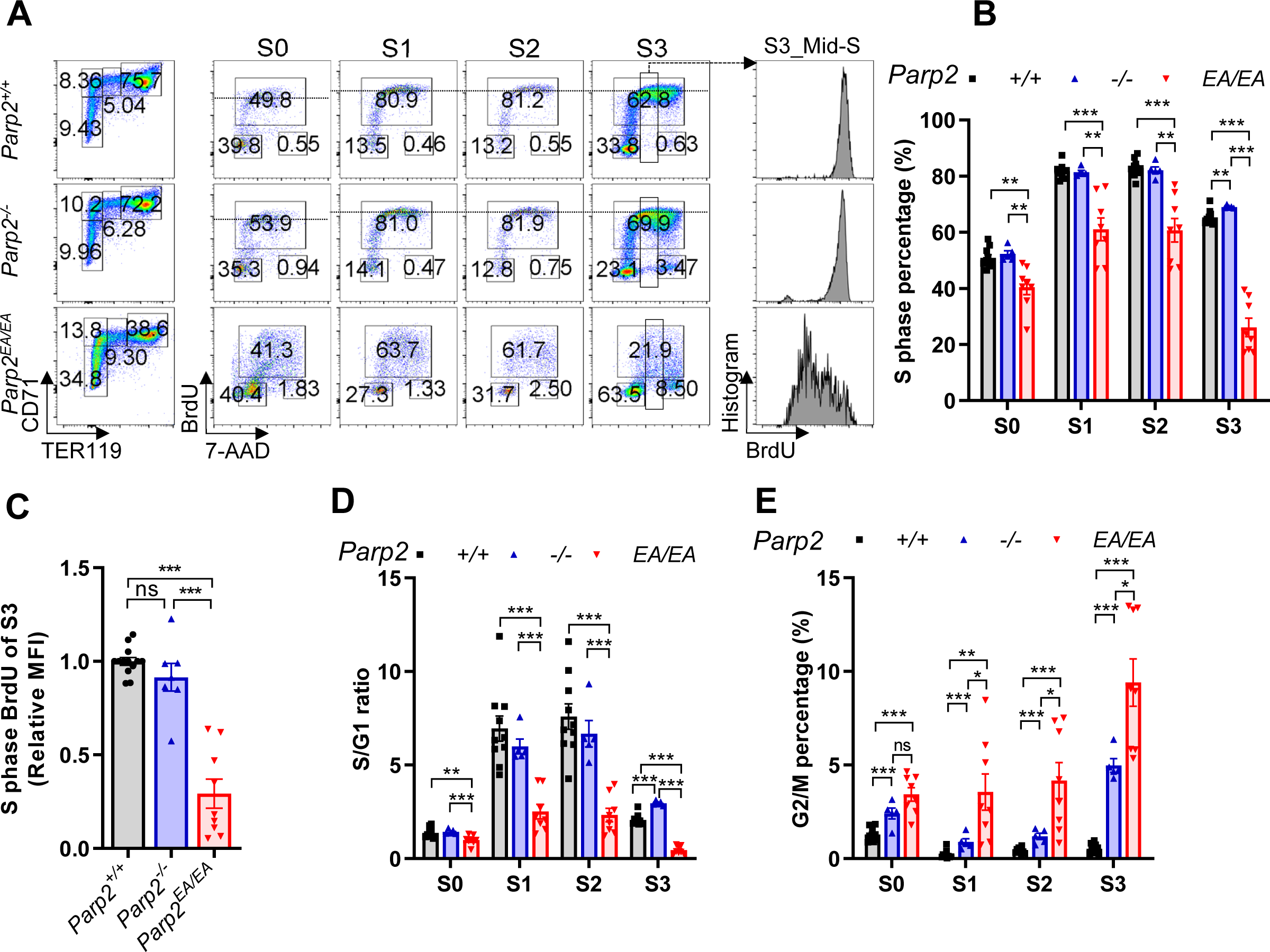
Parp2-E534A blocks cell cycle progression in FL cells. **(A)** Representative flow cytometry analysis of the cell cycle of S0-S3 cells from the FL of E13.5 *Parp2^+/+^, Parp2^-/-^* and *Parp2^EA/EA^* embryos. Dot plots show the percentage of G0/G1, S, and G2/M cells, and the histograms show the distribution of BrdU in mid-7AAD of S3 cells. (B) Quantification of percentage of S phase (BrdU+) cells in S0-S3 fetal livers. Each dot represents an individual embryo of the indicated genotype. (**C**) Relative median fluorescence intensity (MFI) of BrdU in S phase cells among S3 cells from the FL of E13.5 *Parp2^+/+^, Parp2^-/-^*, and *Parp2^EA/EA^* embryos. (**D**) Quantification of the S/G1 ratio and (**E**) the percentage of G2/M cells among S0-S3 cells from the FL of E13.5 *Parp2^+/+^, Parp2^-/-,^* and *Parp2^EA/EA^* embryos. The means ± SEM are shown for all the bar graphs, and an unpaired two-tailed t-test was used to measure the p values. ns, p>0.05; * p<0.05; ** p<0.01; *** p<0.001.

### Loss of Tp53 or Chk2 rescues the lethal anemia in *Parp2^EA/EA^* mice

Tp53 and its upstream kinase CHK2 are major components of the signaling pathways induced by replication stress. Loss of Tp53 and CHK2 have also been linked to high-risk CH that predisposes carriers to myeloid leukemia^7^. Accordingly, we found that the mRNA levels of Tp53 targets (p21, Puma, and Mdm2) are elevated in *Parp2^EA/EA^* FL relative to controls, and this effect is eliminated by Chk2 deletion (fig. S5A-C). Importantly, loss of one or both copies of either the Tp53 or Chk2 gene significantly averts the lethality of *Parp2^EA/EA^*mice, rescues the anemia, and partially restores FL cellularity and erythroblast counts (Fig. 3A-C, and fig. S5D-E). We note that loss of Chk2 (*Parp2^EA/EA^Chk2^-/-^*) consistently achieves a better rescue than loss of Tp53 (*Parp2^EA/EA^Tp53^-/-^*) (Fig. 3A, 3C, and fig. S5D-E), perhaps because defects in the Tp53-mediated G1/S checkpoint (*e.g.*, p21) exacerbate replication stresses in HSC and erythroblasts by allowing damaged cells to initiate replication prematurely^10,37^. Chk2- or Tp53-deficiency largely restored the frequency and intensity of BrdU+ staining and mitotic frequency in *Parp2^EA/EA^* erythroblasts despite the consistent accumulation of some BrdU^low^ cells among *Parp2^EA/EA^Chk2^-/-^* and *Parp2^EA/EA^Tp53^-/-^* S3 cells (Fig. 3D-E and fig. S4A). In addition to the severe anemia, PARPi treatment has been linked with CH^6^ and an elevated risk for treatment-related myeloid leukemia. In this context, loss of CHK2 or TP53 is frequently noted in high-risk CH after genotoxic chemotherapy^6,7^. The ability of Tp53 and Chk2 deficiency to rescue the anemia in *Parp2^EA/EA^* mice suggests that inactivating Parp2 might select for Tp53 and Chk2 loss in HSC, contributing to the leukemia transformation. Nevertheless, the loss of either Chk2 or Tp53 only partially restores erythropoiesis in *Parp2^EA/EA^* mice, indicating that the replication defects persist.

**Figure 3.**
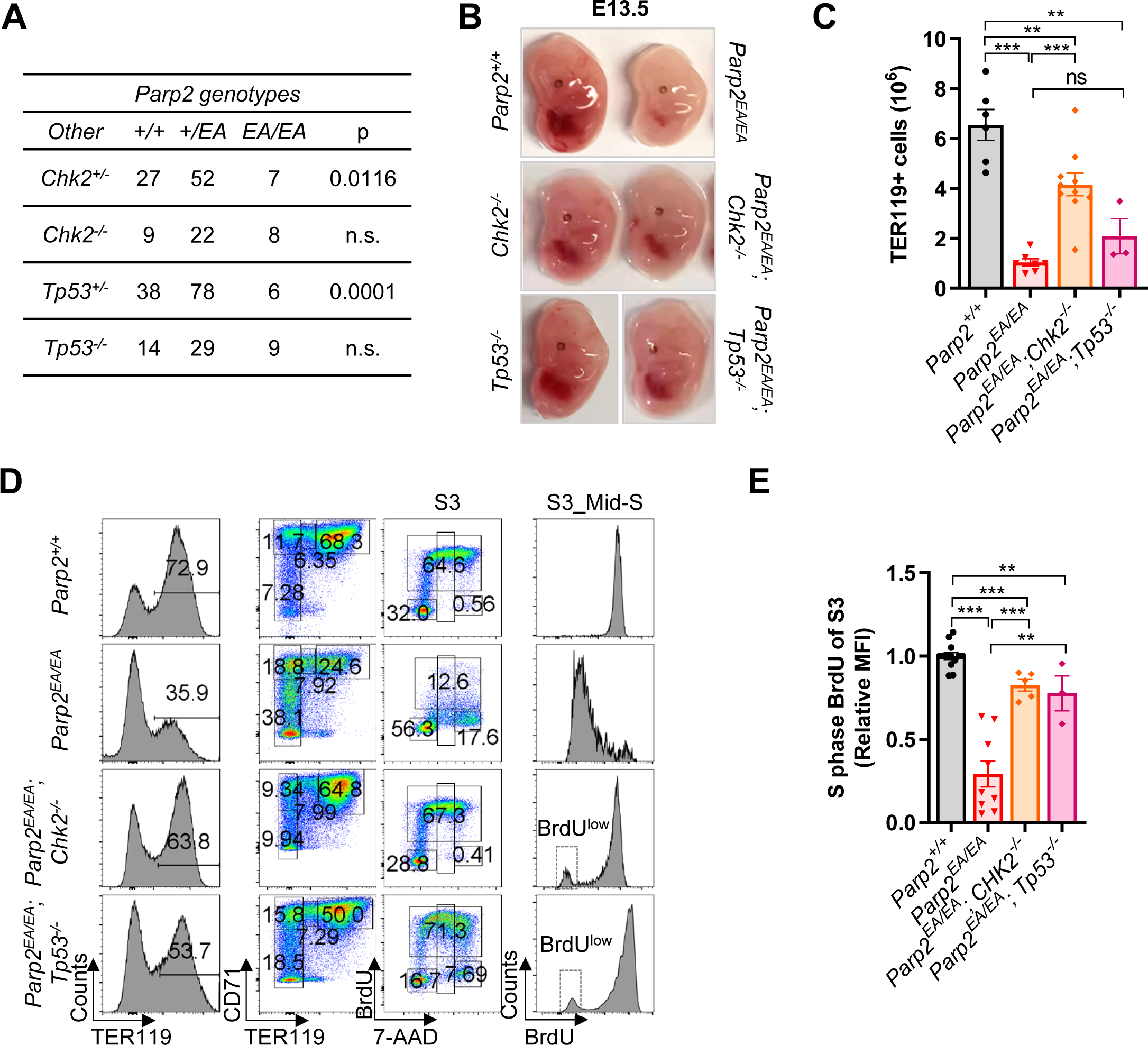
Loss of Tp53 or Chk2 rescues the lethal anemia in *Parp2^EA/EA^* mice. (**A**) Genotype distribution from *Parp2^+/EA^; Chk2^+/-^*intercrossing and *Parp2^+/EA^; p53^+/-^* intercrossing. Chi-squared test. n.s. p>0.05. (**B**) Representative images of E13.5 *Parp2^+/+^*, *Parp2^EA/EA^*, *Chk2^-/-^*, *Parp2^EA/EA^;Chk2^-/-^*, *Tp53^-/-^* and *Parp2^EA/EA^;Tp53^-/-^* embryos. (**C**) Absolute number of TER119^+^ cells from the FL of E13.5 *Parp2^+/+^, Parp2^EA/EA^*, *Parp2^EA/EA^;Chk2^-/-^* and *Parp2^EA/EA^;Tp53^-/-^*embryos. (**D**) Left two panels contain the representative flow cytometry analyses of FL cells from E13.5 *Parp2^+/+^, Parp2^EA/EA^*, *Parp2^EA/EA^; Chk2^-/-^* and *Parp2^EA/EA^; Tp53^-/-^* embryos. The histograms show the percentage of TER119^+^ cells among live cells, and the dot plots show the percentage of S0-S3 cells in the FL. Right two panels: Representative flow cytometry analyses of the cell cycle of S3 cells from the FL of *Parp2^+/+^, Parp2^EA/EA^*, *Parp2^EA/EA^; Chk2^-/-^* and *Parp2^EA/EA^; Tp53^-/-^* embryos. Dot plots show the percentage of G0/G1, S, and G2/M cells and the histograms show the distribution of BrdU in mid-7AAD of S3 cells. (**E**) Relative median fluorescence intensity of BrdU in S phase cells among S3 cells from the FL of E13.5 *Parp2^+/+^, Parp2^EA/EA^*, *Parp2^EA/EA^; Chk2^-/-^* and *Parp2^EA/EA^; Tp53^-/-^* embryos. The means ± SEM are shown for all the bar graphs, and an unpaired two-tailed t-test was used to measure the p values. ns, p>0.05; ** p<0.01; *** p<0.001.

### Inactive PARP2 protein induces replication-dependent genomic instability

Next, we examined the underlying mechanisms responsible for the replication defect of *Parp2^EA/EA^* erythroblasts. Several additional lines of evidence suggest that *Parp2^EA/EA^* erythroblasts accumulate “replication-associated” DNA breaks selectively. First, *Parp2^EA/EA^* FL cells have increased alkaline comet tails, indicative of strand breaks (Fig. 4A). Second, the comet tails of BrdU-negative (none replicating) *Parp2^EA/EA^* cells are not significantly longer than those of wildtype FL cells, indicating that aberrant strand breakage occurs specifically in replicating *Parp2^EA/EA^* cells (Fig. 4B). Third, DNA fiber assays revealed significant fork retardation in *Parp2^EA/EA^* FL cells (Fig. 4C). Consistent with the ultra-fast replication forks in erythroblasts, nucleoside analog labeling duration must be cut to 1/3 of what conventionally used for other cell types (*e.g.*, fibroblasts) to avoid fork merging during the labeling window (fig. S6A). Fourth, BrdU^+^ comet tails, which measure breaks at nascent BrdU+ DNA^27^, are dramatically longer in *Parp2^EA/EA^* FL cells than in either *Parp2^+/+^* or *Parp2^-/-^* control cells (Fig. 4D). Notably, Chk2 deficiency, despite rescuing the anemia, further increased BrdU^+^ comet tails in *Parp2^EA/EA^* FL cells (Fig. 4D), suggesting that CHK2 deficiency allows DNA replication to continue in cells with DNA breaks and exuberates the genomic instability. Correspondingly, *Parp2^EA/EA^* FL cells, especially those in the S or G2 phase, are often positive for phosphorylated histone H2AX (γ-H2AX), a marker of DNA double-strand breaks (DSBs) (Fig. 4E-F). Moreover, the frequency of γ-H2AX staining is even higher in *Parp2^EA/EA^Chk2^-/-^* FL cells (Fig. 4E-F). Together, these observations imply that Parp2^EA^ expression impairs DNA replication by inducing both strand breakage and replication fork collapse. Although the checkpoint defects arising from Tp53 or Chk2 loss can rescue erythropoiesis by enhancing cellular tolerance to DNA breaks, these breaks are not properly repaired in *Parp2^EA/EA^* erythroblasts.

**Figure 4.**
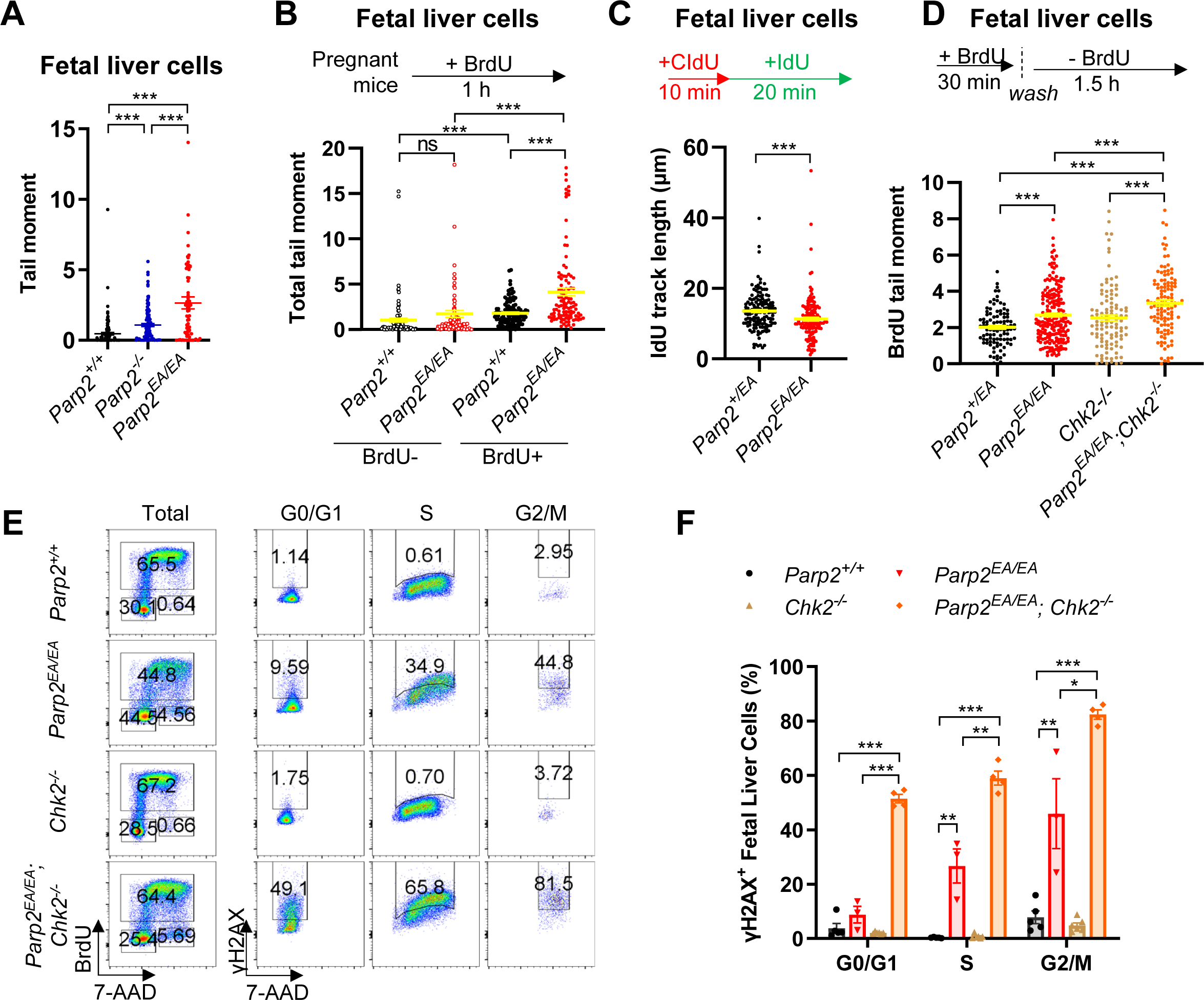
Parp2-E534A induces replication-dependent genomic instability in FL cells. (**A**) Scatter plot of alkaline comet tail moments in total genomic DNA of the FL cells from E13.5 *Parp2^+/+^, Parp2^-/-^* and *Parp2^EA/EA^* embryos. The tail moments of at least 75 individual cells per sample were quantified. (**B**) The alkaline comet tail moment in total genomic DNA of the FL cells from E13.5 *Parp2^+/+^* and *Parp2^EA/EA^* embryos pulse-labeled with BrdU for 1 h. The overall tail moment of BrdU^+^ or BrdU^-^ FL cells was quantified and presented separately. (**C**) The IdU track lengths show the replication fork speed of *in vitro* cultured FL cells from E13.5 *Parp2^+/EA^* and *Parp2^EA/EA^*embryos. Cells were pulse-labeled with 25 µM CldU for 10 min and then with 200 µM IdU for 20 min. At least 100 fibers per sample were quantified. (**D**) BrdU comet tail moment in nascent DNA strands of *in vitro* cultured FL cells from E13.5 *Parp2^+/EA^, Parp2^EA/EA^*, *Chk2^-/-^* and *Parp2^EA/EA^; Chk2^-/-^* embryos. *In vitro,* cultured FL cells were pulse-labeled with 100 µM BrdU for 30 min and subsequent 1.5 h chase. BrdU comet tail moments were scored with anti-BrdU staining to label the nascent DNA strands. (**E**) Representative flow cytometry analysis and (**F**) percentage of γH2AX^+^ cells among the cell cycle of the FL cells from E13.5 *Parp2^+/+^, Parp2^EA/EA^*, *Chk2^-/-^* and *Parp2^EA/EA^; Chk2^-/-^* embryos. For all the bar graphs and scatter dot graphs, the means ± SEM were shown, and an unpaired two-tailed t-test was used to measure the p values. ns, p>0.05; * p<0.05; **, p<0.01; ***p<0.001.

### Inactive PARP2 protein can block Lig1-mediated sealing of 5’p-nicks during DNA replication

We next sought to elucidate the mechanism by which Parp2^EA^ causes replication-associated DNA strand breaks in erythroblasts. Our previous studies of mice expressing kinase-dead forms of ATM, ATR, and DNA-PKcs kinases revealed that the inactive kinase can bind its respective DNA lesion (*e.g.*, ATR to ssDNA and DNA-PKcs to DSBs) and block subsequent repair of that lesion^38-40^. Of note, PARP2 selectively binds to and is consequently activated by 5’p-nicks^41^. During DNA synthesis, 5’p-nicks arise routinely at the junctions between all nascent Okazaki fragments and are typically resolved by DNA Ligase 1 (LIG1) to form the contiguous lagging strand. Accordingly, the PAR level is highest during the S-phase^28^. Given that PARP2 selectively binds 5’p-nicks^41^ and that the inactive Parp2^EA^ displays the trapping behavior of PARPi-treated PARP2 (fig. S1D-I), we tested whether Parp2^EA^ can block LIG1-mediated nick ligation *in vitro*. Indeed, we found that purified WT PARP2 blocks LIG1-mediated nick ligation in the absence, but not the presence, of NAD+ (Fig. 5A). Meanwhile, catalytically inactive Parp2^EA^ blocks nick ligation regardless the presence of NAD+ (Fig. 5A). LIG1-mediated nick ligation in all cell types and is blocked by inactive Parp2, we reasoned that the impact of Parp2^EA^ expression should not be limited to erythroid cells. Indeed, *Parp2^EA/EA^* primary murine embryonic fibroblasts (MEFs) also display Chk2-dependent proliferation defects (Fig. 5B), and fork progression defects (Fig. 5C), and accumulation of BrdU^+^ comet tails (fig. S6B). The flat shape of MEFs also allowed us to measure Lig1 levels at the active fork using multi-color single-molecule localization microscopy (SMLM) and near-neighbor analyses (Fig. 5D). Consistent with the replication defects, productive DNA synthesis measured by chromatin-associated EdU, Pcna, and Lig1 all decreased in *Parp2^EA/EA^*primary MEFs (fig. S6C-E). Moreover, the association of Lig1 to replicative chromatin (i.e., marked by Pcna or EdU) is clearly reduced in *Parp2^EA/EA^*but not in *Parp2^-/-^* primary MEFs (Fig. 5E and fig. S6F). In all cases, the association of chromatin-bound Lig1 with Pcna or EdU is markedly higher than expected from a random distribution, suggesting the specificity of the association (fig. S6G-H). Together, these data support a model in which inactive Parp2 binds 5’p-nicks and interferes with Lig1-mediated Okazaki fragment maturation.

**Figure 5.**
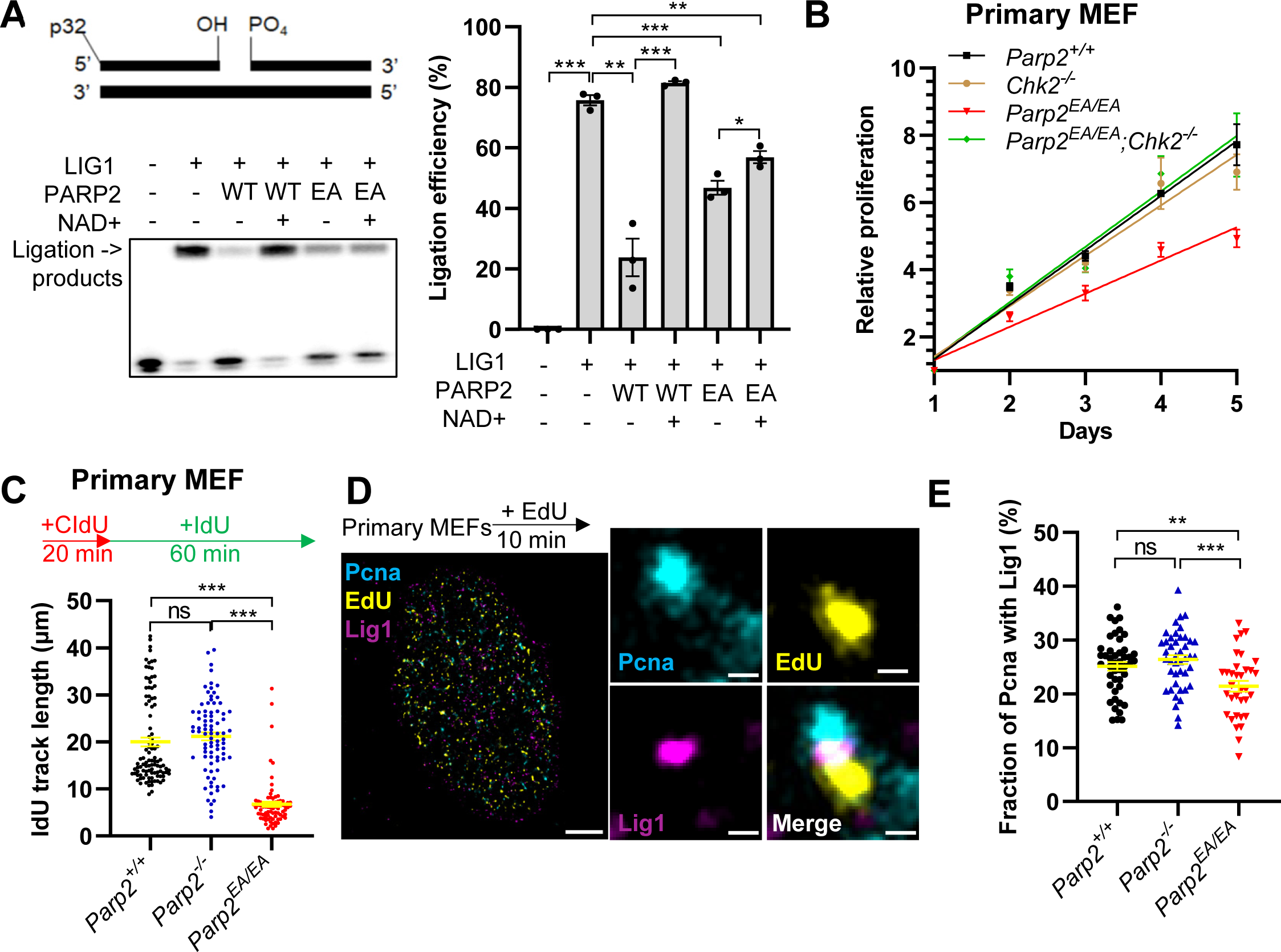
Inactive PARP2 impairs DNA replication by blocking LIG1-mediated nick ligation. (**A**) The representative gel image (left panel) and quantification (right panel) of *in vitro* LIG1 mediated nick-ligation in the presence of purified PARP2 WT or E545A protein with or without NAD+. About 0.5 pmol of nicked DNA substrate was pre-incubated with or without 2 pmol of PARP2 before exposure to 0.25 pmol of DNA LIG1. The average and standard errors of the three reactions were shown. (**B**) Relative proliferation of primary MEFs derived from E13.5 *Parp2^+/+^, Parp2^EA/EA^*, *Chk2^-/-^* and *Parp2^EA/EA^; Chk2^-/-^* embryos, measured with MTT assay. (**C**) IdU track lengths show the replication fork speed in *Parp2^+/+^*, *Parp2^-/-^,* and *Parp2^EA/EA^* primary MEFs. Cells were pulse-labeled with 25 µM CldU for 20 min and then with 200 µM IdU for 1 h. At least 100 fibers per sample were quantified. (**D**) Representative SMLM images of a *Parp2^+/+^* primary MEF nucleus pulse-labeled with EdU (yellow) for 10 min followed by immunostaining for Pcna (blue), and LIG1 (magenta). Scale bar: Whole nuclei = 2500 nm and inset = 100 nm. (**E**) Fraction of Pcna clusters associated with Lig1 within a 40 nm radius in *Parp2^+/+^, Parp2^-/-,^* and *Parp2^EA/EA^* primary MEFs. Individual data points represent values from a single nucleus determined by DBSCAN/NND analysis. n>30 nuclei were analyzed in three independent biological replicates. For all the scatter dot graphs and bar graphs, the means ± SEM were shown, and an unpaired two-tailed t-test was used to measure the p values. ns, p>0.05; * p<0.05; **, p<0.01; ***p<0.001.

If so, *Parp2^EA/EA^* and Lig1-deficient mice should have similar phenotypes. While mice with a C-terminal deletion of the *Lig1* gene retaining the catalytic exons (*Lig1^Δ/Δ^)* die of anemia before birth^30^, this mouse strain is no longer available. Therefore, we generated a new Lig1-null strain in which Lig1 exons 3-5, which lie upstream of coding sequences for the DNA binding and adenylation domains, have been removed (fig. S7A). As expected, *Lig1^-/-^* cells express no detectable Lig1 protein (fig. S7B), the E13.5 *Lig1^-/-^* embryos are pale (Fig. 6A) and display profound erythropoiesis defects. Interestingly, however, the erythroid and B cell defects of *Lig1^-/-^* embryos consistently less severe than those of *Parp2^EA/EA^* embryos (Fig. 6B-C and fig. S7C), suggesting other DNA ligases might partially compensate for the loss of Lig1 in *Lig1^-/-^* cells, and that inactive Parp2 would also block Okazaki fragment maturation by these ligases in *Parp2^EA/EA^*cells.

**Figure 6.**
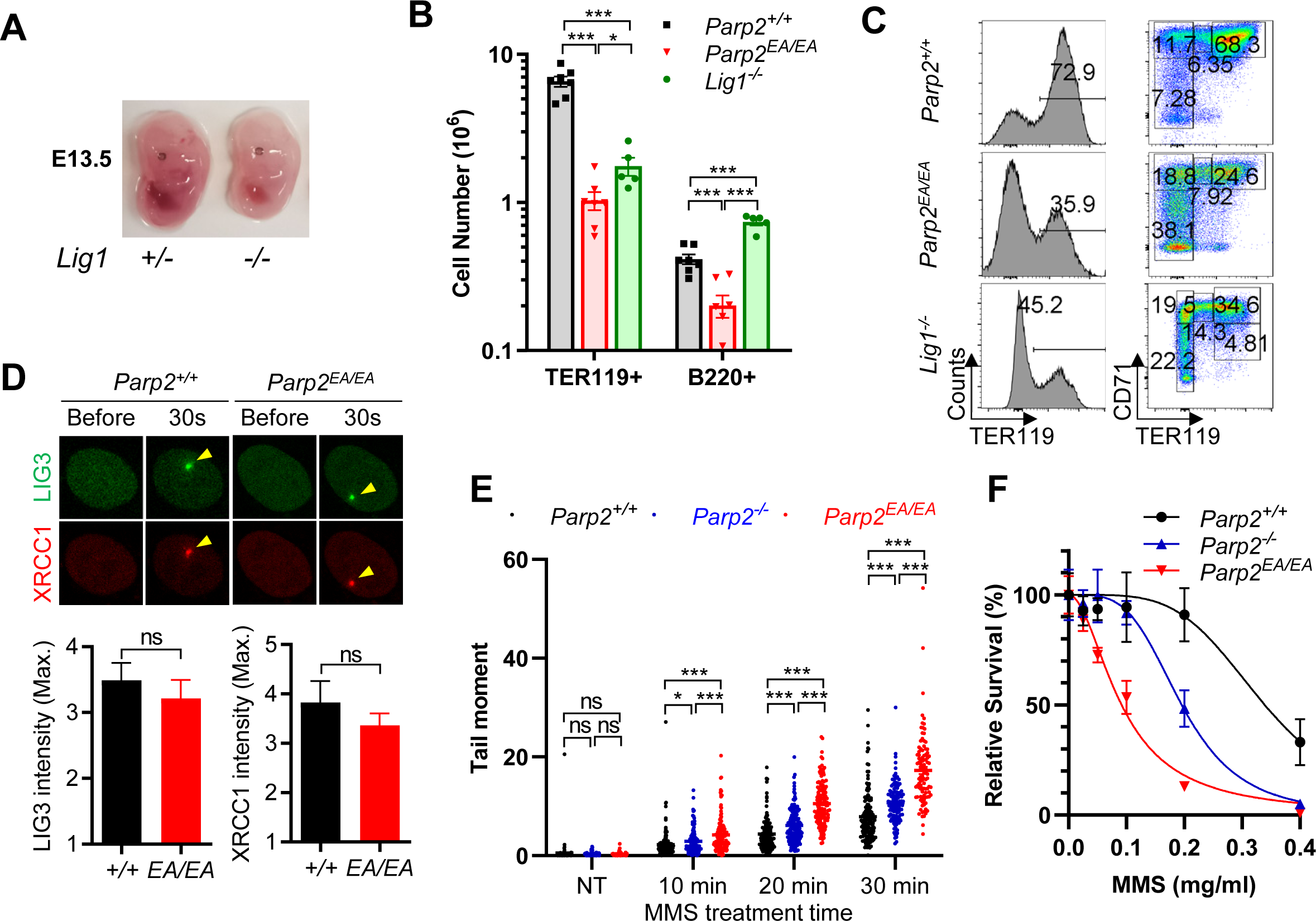
Inactive PARP2 impedes LIG3-mediated DNA ligation. (**A**) Representative images of *Lig1^+/-^* and *Lig1^-/-^* embryos at E13.5. (**B**) The absolute number of TER119^+^ and B220^+^ cells in the FL of E13.5 *Parp2^+/+^, Parp2^EA/EA,^* and *Lig1^-/-^* embryos. (**C**) Representative flow cytometry analyses of FL cells from E13.5 *Parp2^+/+^, Parp2^EA/EA,^* and *Lig1^-/-^* embryos. The histograms show the percentage of TER119^+^ cells among live cells, and the dot plots show the percentage of S0-S3 RBC in the FL. (**D**) Representative images (upper panel) and the maximal relative intensity (lower panel) of EGFP-LIG3 and mRFP-XRCC1 at DNA damage sites induced by 405 nm micro-irradiation. (**E**) Scatter plot of alkaline comet tail moments in total genomic DNA of *Parp2^+/+^, Parp2^-/-^* and *Parp2^EA/EA^* iMEFs treated with 0.1 mg/ml MMS for 10, 20, and 30 min. (**F**) The sensitivity assay of *Parp2^+/+^, Parp2^-/-,^* and *Parp2^EA/EA^*iMEFs to MMS. Cells were treated with indicated concentrations of MMS for 30 min, followed by incubation in a drug-free medium for 5 days. For all the scatter dot graphs and bar graphs, the means ± SEM were shown, and an unpaired two-tailed t-test was used to measure the p values. ns, p>0.05; * p<0.05; **, p<0.01; ***p<0.001.

### Inactive PARP2 delays Lig3-mediated repair without compromising Lig3 recruitment

Although Okazaki fragment ligation is primarily mediated by LIG1, LIG3 has been proposed as the backup mechanism to ensure the continuity of lagging strand synthesis. Accordingly, Lig1 is not essential for DNA replication in most mammalian cell types tested^29^, except during erythropoiesis^30^. The *LIG3* gene encodes both a non-essential nuclear isoform and an essential mitochondrial isoform^42,43^. Nuclear LIG3 forms an obligatory complex with XRCC1, which, by virtue of its PAR-binding BRCT1 domain, can recruit LIG3, other BER proteins (*e.g.*, Polβ) to DNA breaks^44^. For example, during BER, damaged DNA bases, such as those induced by alkylating agents (*e.g.*, methyl methanesulfonate/MMS), are removed by glycosylases to generate abasic sites, which are in turn converted into a 5’p-nicks by the AP endonuclease. Upon binding these nicks, enzymatically-activated PARP1 and PARP2 generate PAR chains that recruit the XRCC1-LIG3 complex for nick ligation^44^. Consistent with the dominant role of PARP1 in alkylating agent-induced PARylation^8^, the PAR-dependent recruitment of XRCC1 and LIG3 to micro-irradiation sites wa unaffected in immortalized MEFs (iMEFs) from *Parp2^EA/EA^* embryos (Fig. 6D and fig. S7D-F). Nonetheless, higher levels of MMS-induced breaks measured by comet tails were observed in *Parp2^EA/EA^*iMEFs than in *Parp2^-/-^* or *Parp2^+/+^* controls, suggesting that the presence of Parp2^EA^ delays end ligation (Fig. 6E). Correspondingly, *Parp2^EA/EA^* iMEFs are more sensitive to MMS than *Parp2^-/-^* and *Parp2^+/+^*cells (Fig. 6F). *Parp2^EA/EA^* iMEFs are also hypersensitive to the TopoI inhibitor camptothecin (CPT), which generates 5’p-nick intermediates (fig. S8A)^45^. Meanwhile, neither *Parp2^EA/EA^* nor *Parp2^-/-^* iMEFs are sensitive to the TopoII inhibitor etoposide or to ionizing radiation, both of which primarily induce DSBs (fig. S8B-C). *Parp2^EA/EA^* and *Parp2^-/-^* iMEFs are equally, and only moderately, sensitive to agents that retard replication fork progression, such as hydroxyurea (HU), aphidicolin, and the crosslinking agent mitomycin C (MMC) (fig. S8D-F). Together, these data indicate that inactive Parp2 delays Lig3-mediated repair without affecting the PAR-dependent recruitment of Xrcc1-Lig3 to 5’p-nicks. A similar effect has been described for inactive DNA-PKcs protein, which allosterically blocks LIG4-mediated repair without impairing KU-dependent recruitment of LIG4 to laser-induced DNA damage^40,46^.

### PARP2 is significantly activated by normal DNA replication

The recruitment of Lig3 to micro-irradiation sites in *Parp2^EA/EA^*cells is consistent with a dominant role for PARP1-mediated PARylation during the DNA damage response. Yet, why is the presence of inactive Parp2 sufficient to block DNA replication in *Parp2^EA/EA^* erythroblasts despite the expression of wild-type Parp1 protein? Of note, PARP1/2-mediated PARylation is usually studied upon acute external DNA damage (*e.g*., MMS and H_2_O_2_) that generates diverse DNA lesions. In contrast, erythropoiesis occurs in the absence of overt external genotoxic stress. Undisturbed DNA replication in erythroblasts consistently generates 5’p-nicks between Okazaki fragments. Therefore, to assess the relative activities of Parp1 and Parp2 in undamaged cells, we compared endogenous auto-PARylation of the Parp1 and Parp2 in wildtype E13.5 embryos. Not surprisingly, comparable levels of PARylated Parp1 were observed in the fetal body (without fetal brain and FL) and FL (>70% erythroblasts) from the same mice (Fig. 7A). In contrast, PARylated Parp2 levels, though barely detectable in the fetal body, are dramatically higher in FL cells (Fig. 7A). Levels of Chk2, expressed during the S/G2 phases, are also higher in FL, suggesting that Parp2 PARylation is occurring in replicating cells (Fig. 7A). Notably, PARylated Parp2 is absent in both *Parp2^-/-^* and *Parp2^EA/EA^* FLs (Fig. 7B), indicating that Parp1 cannot trans-PARylate Parp2. Phosphorylated Chk1, a marker for replication stress^47^, and caspase-cleaved Parp1 and Parp2 were only detected in *Parp2^EA/EA^* FL, suggesting that Parp2^EA^-expressing FL cells experience replication stress and undergo apoptosis (Fig. 7B and fig. S2E). Also, the replication-associated auto-PARylation of Parp2 is not restricted to erythroblasts, as PARylated forms of both Parp1 and Parp2 are also abundant in untreated iMEFs. As expected, MMS treatment induces hyper-PARylation of Parp1 (Fig. 7C). In contrast, however, MMS reduces Parp2 PARylation, possibly by blocking replication fork progression and thus the formation of Okazaki-flanking 5’p-nicks (Fig. 7C). Thus, although PARP1 is the predominant mediator of PARylation during the DNA damage response, our results reveal an unexpected role for PARP2 in normal DNA replication that is consistent with its 5’p-nick selectivity and the need to resolve the 5’p-nicks that arise during lagging strand synthesis.

**Figure 7.**
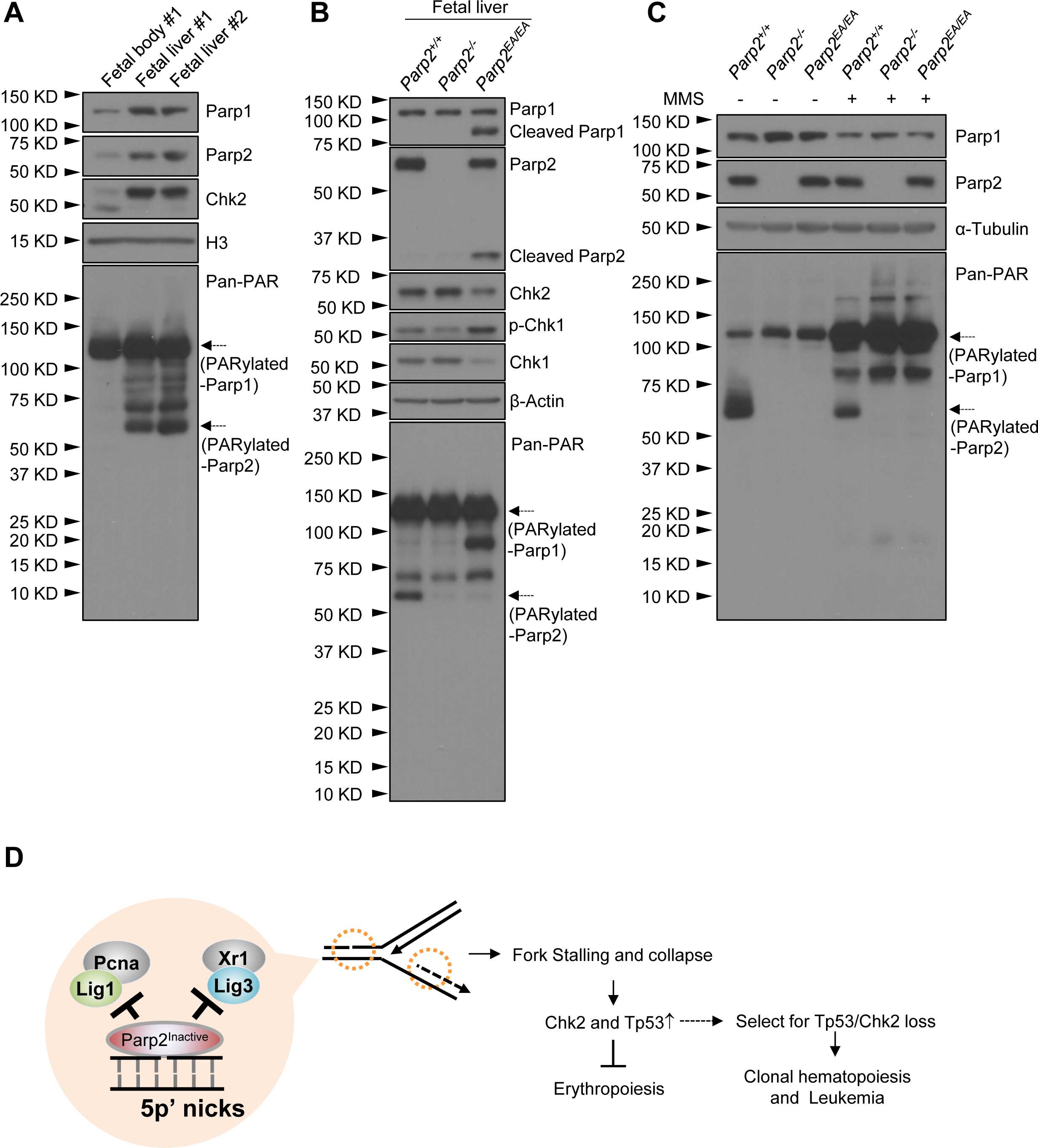
Parp2 PARylation is enriched in the fetal liver and reduced by MMS. (**A**) Western blot to analyze the endogenous PARylation levels of Parp1 and Parp2 in the fetal body and FL of normal E13.5 embryos. Fetal body #1 and FL #1 were obtained from the same embryo #1. Embryo #1 and #2 were littermates. (**B**) Western blot to analyze the endogenous PARylation levels of Parp1 and Parp2 in FLs from E13.5 *Parp2^+/+^, Parp2^-/-,^*and *Parp2^EA/EA^* embryos. (**C**) Western blot to analyze the endogenous and DNA damage-dependent PARylation levels of Parp1 and Parp2 in *Parp2^+/+^, Parp2^-/-,^*and *Parp2^EA/EA^* iMEFs. DNA damage-dependent PARylation was induced by treating cells with 0.1 mg/ml MMS for 1 h before harvest. (**D**) The working model shows that inactive PARP2 binds to DNA nicks at lagging strands or in repair intermediates to block Lig1 and Lig3 mediated end-ligation and causes replication fork stalling and collapse, Chk2 and Tp53 activation and erythropoiesis failure.

## Discussion

Collectively, our data show that catalytically inactive Parp2 (Parp2^EA^) selectively binds to and is allosterically retained at 5’p-nicks, including those formed during normal DNA replication, and that its presence blocks Lig1 and Lig3-mediated nick ligation required for Okazaki fragment maturation. In fast-replicating erythroblasts that rely on the timely processing of Okazaki fragment, the presence of inactive Parp2^EA^ triggers frequent replication fork collapse, DNA strand breakage, and eventually Tp53- and Chk2-dependent checkpoint activation and apoptotic cell death, resulting in lethal anemia (Fig. 7D). Moreover, we propose that the preferential impact of Parp2^EA^ on erythropoiesis and erythroblast can be explained by the DNA lesion selectivity of PARP2 to the 5’p-nicks, which consistently arise at the nascent lagging strands during replication. Indeed, in contrast to external DNA damage (*e.g.*, alkylating agents) that preferentially activates PARP1, undisturbed DNA replication with consistent presence of 5’p-nick activates PARP2 robustly (Fig. 7A-C). The presence of inactive PARP2, but not active PARP2, or the absence of PARP2 physically blocks nick ligation by Lig1 (Fig.5) and Lig3 (Fig.6). In this respect, we show that the behavior of Parp2^EA^ resembles that of PARPi treated wild-type PARP2^14,15^, including allosteric retention and persistence. Thus, we propose that the unexpected severe anemia associated with PARPi maintenance therapy arises through a similar molecular mechanism: namely, that PARPi delays the release of PARP2 from the 5’p-nicks on the lagging strands, where they block nick ligation and impairs Okazaki fragment maturation, detrimental for erythroblasts with fast forks. We note that the physical block to nick ligation imposed by inactive PARP2 is not limited to lagging strands and can also occur at other 5’p-nicks, such as those that arose during BER (Fig. 6E-F and 7D), further exacerbating the replication defects. Finally, we show that loss of TP53 or CHK2 restores the erythropoiesis and rescues the embryonic lethality of *Parp2^EA/EA^* FLs, revealing a direct connection between PARP2 inhibition and clonal hematopoiesis, especially in the context of TP53 or CHK2 loss. Given loss of TP53 and CHK2 are often associated with high-risk CH upon genotoxic therapy, this finding provides a molecular explanation for unexpected severe anemia and treatment-related myeloid leukemia associated with PARPi therapy.

While our results indicate that inactive Parp2 can potentially block DNA replication in multiple cell types, including fibroblasts and B lymphocytes, why is erythropoiesis especially vulnerable in *Parp2^EA/EA^* mice? We propose that this tissue-specificity reflects the exceptionally high proliferation demand during erythropoiesis and the unique features of erythroblast replication. Despite their small size, red blood cells account for ∼50% of the blood by volume, while all white blood cells combined represent 1%. Erythropoiesis can be briefly divided into progenitor and terminal differentiation phases^35^. At the transition to the terminal differentiation phase, S1 erythroblasts undergo a differentiation-coupled replication program characterized by not only very short gap phases (thus high BrdU+% in pulse label experiments) but also ultra-fast fork speeds coupled with progressive loss of DNA methylation that is typically associated with replication^31,33,35^. Indeed, BrdU+% and BrdU levels per cell are highest in S1 erythroblasts compared to S0 progenitors or FL B cells (Fig. 2A and fig. S4). The fork speed in erythroblasts is also higher than in primary fibroblasts (fig. S6A). Consistent with a need for highly efficient Okazaki fragment maturation in erythroblasts, DNA ligase 1 null mice also succumbed to anemia due to FL erythropoiesis failture^30,48^, which is strikingly similar to that observed in *Parp2^EA/EA^* embryos. In contrast, cancer cells or early embryonic tissues often achieve fast growth by increasing CDK activities, which leads to the simultaneous initiation of a greater number of replication origins^36^. We speculate that this difference in replication fork speed might account for the differential sensitivity of FL B cells vs. erythroblasts to inactive Parp2 and why the Parp2 inhibition is particularly detrimental for erythropoiesis. Together, these data identified replication fork speed as a new predictor for PARPi sensitivity.

Consistent with physical blocking by the inactive PARP2 proteins, the severity of the erythropoietic defect *in vivo* correlates with the dose of inactive Parp2^EA^ protein but not the enzymatic activity of Parp2. As such, all *Parp2^EA/EA^* embryos die of anemia, *Parp2^-/EA^* pups are born alive, albeit under-representative, and display severe anemia at a young age (4-6 weeks), and *Parp2 ^-/-^* mice are born healthy at the Mendelian ratio and develop moderate anemia later (>90 days/12 weeks). If the anemias of *Parp2^EA/EA^* and *Parp2^-/EA^*mice are caused by inactive Parp2^EA^ protein that blocks 5’p-nick ligation, how can we explain the mild anemia of adult *Parp2^-/-^* mice? We noted that the exchange and release of PARP1 is significantly slower than PARP2 in the FRAP experiments (fig. S9A-B, t_1/2_ ≈ 2.4s for PARP2 and 7.7s for PARP1)^15,16^. In this regard, PARP2 binds to 5’p-nick with its sole WGR domain, while PARP1 has three ZnF domains, the BRCT domain and the WGR domain, all of which bind to DNA. Collectively, PARP1 has more extensive DNA binding footprints, which might contribute to the slower release. Given fast fork progression in erythroblasts likely sensitive to timely ligation of 5’p-nicks, we propose that in the absence of PARP2, PARP1 will be recruited to the 5’p-nicks and the relatively slow release of PARP1 might impede 5’p-nick ligation and compromise erythropoiesis, thus contributing to mild anemia in adult *Parp2^-/-^*mice^10^. Alternatively, the presence of active PARP2 might facilitate the repair and reduce PARP1-trapping by promoting the recruitment of XRCC1-Lig3 and subsquent nick-ligation by providing additional PAR signal through auto-PARylation. A similar mechanism might explain the mild T-cell development defects in *Parp2^-/-^* but not *Parp1^-/-^* mice. While we were unaware of any direct measurement of replication fork speed in developing T cells, germinal center B lymphocytes activated via the help of follicular T cells are also known to have fast fork speed^49^.

How does enzymatically inactive Parp2^EA^ block DNA replication despite the presence and relative abundance of Parp1? Although PARP1 and PARP2 are released from DNA ends independent of PARylation, extensive auto-PARylation prevents PARP1 from binding back to DNA^50^. Consequently, whenever auto-PARylated PARP1 is released from DNA ends, inactive PARP2, despite its lower abundance, could gradually occupy and hinder the ligation of more 5’p-nicks through allosteric retention^14,15^. This effect, as well as the preferential binding of 5’p-nicks by PARP2 and the continual formation of 5’p-nicks between newly synthesized Okazaki fragments during replication, likely explains how inactive Parp2 can impede erythropoiesis in the presence of abundant wildtype Parp1. Nonetheless, it is also possible that PARP2 is actively recruited to replicate DNA by yet unknown mechanisms, especially given the dramatic induction of Parp2 auto-PARylation accompanies normal DNA replication despite its low abundance (Fig. 7). In this context, we previously showed that PARP1 and PARP1 activity could promote the local enrichment of PARP1^15^. Moreover, PARP2 has an unstructured N-terminal region that, while dispensable for break-induced PARP2 activation, can also bind to DNA or even PAR^51,52^. In any case, our findings do not exclude a significant role for Parp1 as a sensor for unligated Okazaki fragments^28^. But, given the importance of inactive PARP1 in cancer therapy, sparing PARP1 would not be a fruitful strategy to minimize the hematological toxicity of PARPi. Consistent with a broader impact of PARP1 inactivation, we recently described the phenotypes of mice expressing an inactive Parp1 with a comparable catalytic site mutation (Parp1-E988A)^21^. Whereas Parp1-null mice develop normally^9^, heterozygous *Parp1^+/E988A^* mice expressing both wildtype and inactive Parp1 die before embryonic day E9.5, indicating that Parp1-E988A expression disrupts embryonic development in a dominant-negative and much more severe fashion^21^ than the relatively specific lethal anemia of *Parp2^EA/EA^* embryos and *Parp2^-/EA^* pups described here. In this context, we note that *Parp2^+/EA^*mice with one wildtype Parp2 allele have no measurable phenotypes (Fig. 1A and 1F, and fig. S2B-C). Thus, the earlier embryonic lethality of *Parp1^+/E988A^* mice (before E9.5) indicates that inactive Parp1 impacts the viability of a broader spectrum of cell types at an earlier stage without the preferential effects of inactive PARP2 on erythropoiesis.

As noted above, the behavior of inactive Parp2^EA^ in our mouse models resembles that of wildtype PARP2 in patients undergoing PARPi therapy. In either setting, the PARylation activity of PARP2 is impaired, and the inactive protein (Parp2^EA^ or PARPi-bound PARP2) displays the characteristic features of “PARP trapping”, such as prolonged retention in the chromatin fraction and at laser-induced DNA damage sites^12,13^, and slower exchange/release^14,15^. Since erythropoiesis is especially sensitive to inactive Parp2 in mice, it is likely that inhibition of human PARP2 function also contributes to the anemia that arises during PARPi maintenance therapy. At the molecular level, expression of inactive PARP1/2 and exposure to PARPi both cause dramatic delays in the exchange of PARP2 on DNA ends without significantly affecting PARP1 exchange^14-17^. Moreover, our FRAP studies show that FDA-approved PARPi agents with potent hematological toxicity (i.e., niraparib and talazoparib) extend the t_1/2_ of DNA binding by wildtype PARP2 to a greater extent (nearly 20 seconds) than olaparib (∼10 seconds), a PARPi with low hematological toxicity^15,16^. These observations argue that the anemic toxicities arising in PARPi-treated patients are partly caused by the inhibition of PARP2 activity. In light of existing evidence that the loss of PARP1, but not PARP2, confers significant PARPi resistance in cancer cells and gRNAs targeting PARP1, but not PARP2 are enriched in CRISPR screen with PARPi^15,16,53,54^, our data suggest that by avoiding PARP2 allosteric trapping, new PARPi can minimize hematological toxicity without compromising its therapeutic efficacy. Indeed, a newly developed PARP1-selective inhibitor was recently shown to elicit milder hematological toxicity in pre-clinical models^55^.

In summary, our data supports a model in which, upon selectively binding to 5’p-nicks, inactive Parp2 disrupts Okazaki fragment maturation by blocking Lig1- and Lig3-mediated nick ligation on the lagging strand of replicating DNA. Although inactive Parp2 affects DNA synthesis in multiple cell types, terminal-differentiating erythroblasts, which rely on an ultra-fast replication fork progression, are especially sensitive to its effects. In particular, DNA strand breaks and collapsed replication forks accumulate in these cells, and the resulting genome instability induces Tp53- and Chk2-dependent erythropoiesis failure and ultimately lethal anemia (Fig. 7D). By selecting for the loss of Tp53 or Chk2 to support erythropoiesis, inactivating Parp2 promotes clonal hematopoiesis, especially those with high risk for leukemia transformation. Our study also reveals a curious similarity between the deletion versus inactivation of DNA damage response proteins, such as PARP1^21^, PARP2, and the three DNA damage response kinases (ATM^39^, ATR^38^, and DNA-PKcs^40^) governing cellular responses to genotoxic stress. In each case, the expression of a catalytically inactive polypeptide yields a more severe phenotype than the deletion of the protein, as the inactive polypeptide can be “trapped” at its cognate DNA lesion, hindering DNA repair by other proteins or pathways^56^. To fully understand the therapeutic impacts of small molecular inhibitors thus requires us to go beyond deletion and adventures into catalytic inactivation models. Finally, by providing a mechanistic explanation for the severe and unforeseen hematological toxicity associated with current PARP inhibitors, our study offers insights into the nuanced interplay of and the functional difference between PARP1 and PARP2 in DNA repair and replication that can be explained by their DNA lesion specificity and pave the way for refining the therapeutic windows of PARPi and other DNA damage inhibitors.

## Acknowledgments

We greatly appreciate discussions and suggestions from Drs Yonghao Yu, Richard Baer, and members of the Zha lab. We thank Dr. Victor Lin at the HICCC transgenic core for his assistance with ES cell injection. The work is supported by NIH CA226852, CA271595, CA174653, CA293675 to SZ; NIH CA275184 to SZ, ER; NIH P30CA013696 to HICCC; and NIH GM057479 and CA276837 to AT; and NIH NS37956 and CA21765 to PM. PM’s lab is also supported by American Lebanese and Syrian Associated Charities of St. Jude Children’s Research Hospital.

## Author contributions

X.L. generated and characterized the Parp2 inactive mouse models designed and performed most experiments. D.G. and E.R. provided the instruments, training, and analytic tools necessary for single-molecule imaging. A.V. performed the BrdU comet analyses in MEFs. K.L. helped with western, comet assay, DNA fiber assay, and quantitative live cell imaging analyses. D.M. advised on DNA fiber analyses and comet analyses. S.K.B. and A.E.T. purified the PARP2 and LIG1 proteins, performed the *in vitro* LIG1 mediated ligation analyses, and provided the LIG1 antibody used in western blot and single-molecule imaging. H.R.R. and P.M. generated the Lig1 conditional mouse model. W.G. advised us on the Tp53 target RT-PCR analyses. F.D. generated the Parp2 null and conditional null mouse models. B.J.L. and S.G. helped with mouse colony and genotyping. X.L. and S.Z. conceived the project and wrote the manuscript with input from other co-authors.

## Data and materials availability

All data and materials presented here are available in the public database or upon request.

## Supplementary Figure Legends

**Figure S1.**
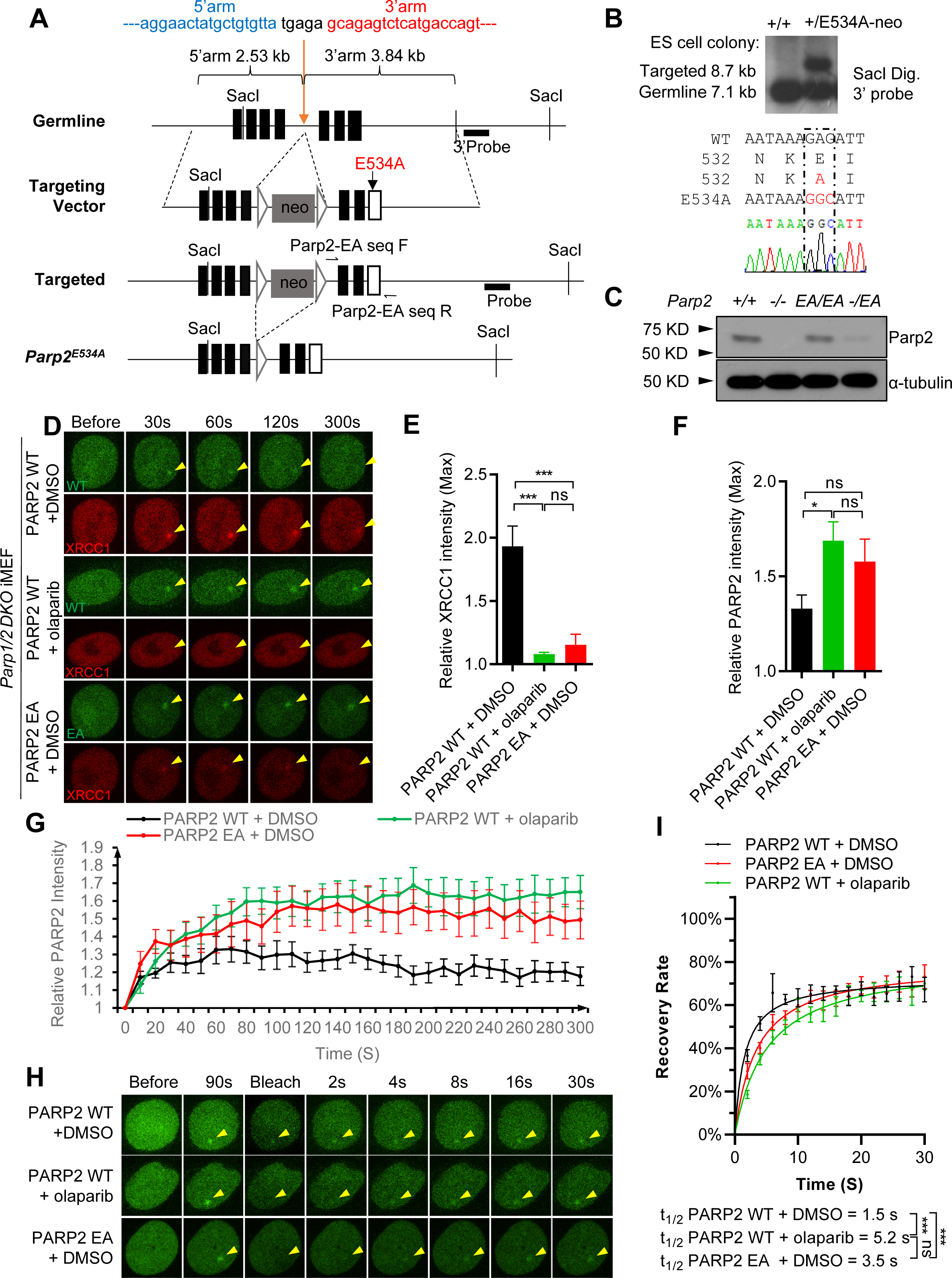
Generation and validation of Parp2-E534A mouse model. **(A)** Targeting scheme of the *Parp2^E534A^* allele. The exons are solid-filled black boxes, except the targeted exon containing the E534A mutation is an open box. The FRT sites are shown as blank triangles. The probe for southern blotting is marked as a thick black line. The primers (Parp2-EA seq F and Parp2-EA seq R) are marked with arrows for amplifying and sequencing the E534A mutation. The locations of SacI restriction sites for genome digestion are marked. **(B)** Southern blot analyses and the sequencing results of successfully targeted ES cells. Genomic DNA was digested with SacI restriction enzyme and detected by southern blot with the probe. (**C**) Western blot confirmed the expression level of Parp2 or Parp2^EA^ in primary MEFs from E13.5 *Parp2^+/+^, Parp2^-/-^*, *Parp2^EA/EA,^* and *Parp2^-/EA^* embryos. (**D)** Representative images of EGFP-PARP2 and mRFP-XRCC1 foci formation. (**E**) The maximal relative intensity of mRFP-XRCC1. (**F**) The maximal relative intensity of EGFP-PARP2 and (**G**) the relative intensity kinetics of EGFP-PARP2 at DNA damage sites induced by 405 nm micro-irradiation in *Parp1/2 DKO* iMEF. The means ± SEM are shown for all the bar graphs, and an unpaired two-tailed t-test was used to measure the p values. *** p<0.001.

**Figure S2.**
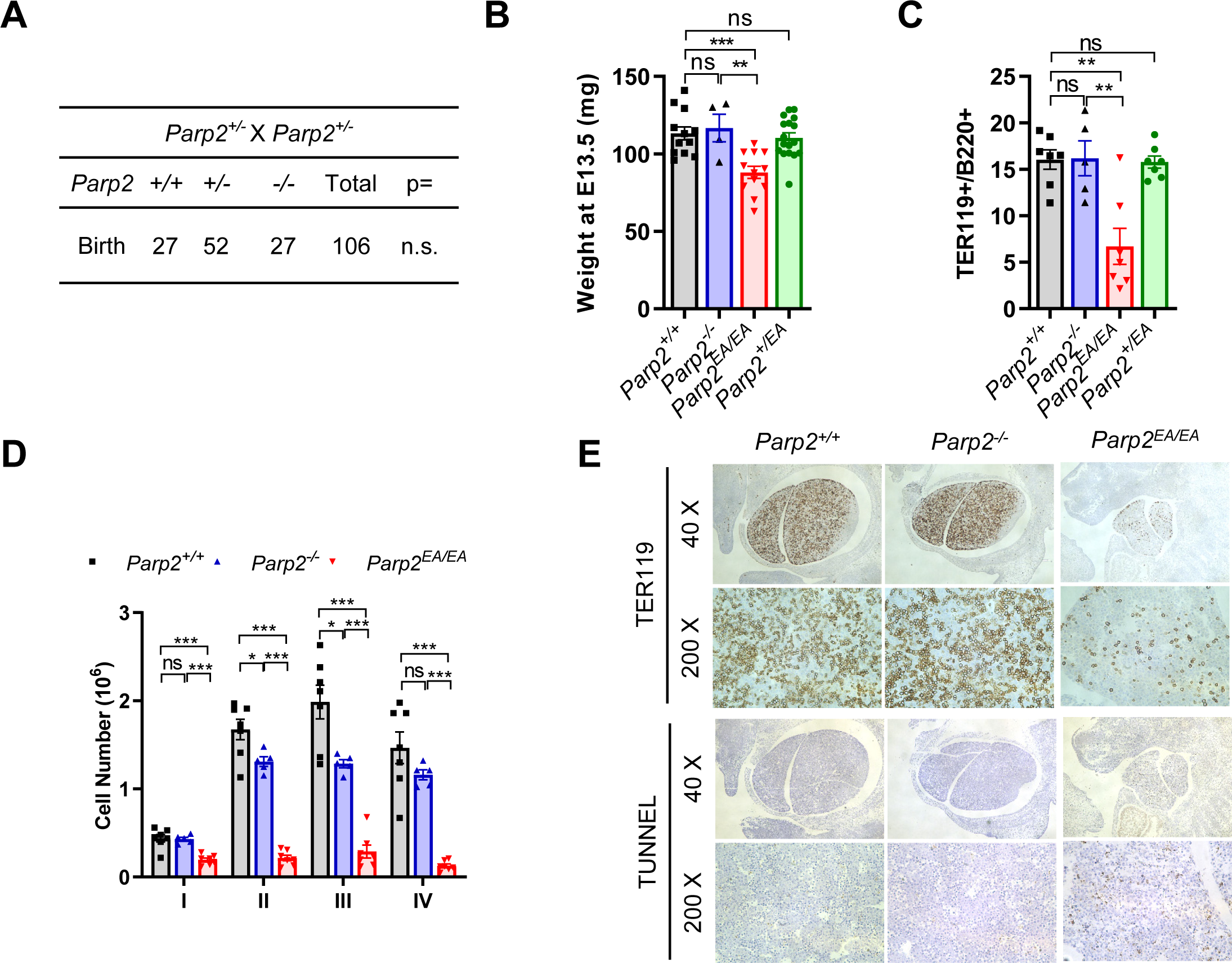
Hematopoiesis analysis of *Parp2^EA/EA^* FL. (**A**) Genotype distribution from *Parp2^+/-^* intercrossing. Chi-squared test. n.s. p>0.05. (**B**) Weight of E13.5 *Parp2^+/+^, Parp2^-/-^*, *Parp2^EA/EA^* and *Parp2^+/EA^* embryos. (**C**) Ratio of TER119^+^ and B220^+^ cells in FLs from E13.5 *Parp2^+/+^, Parp2^-/-^*, *Parp2^EA/EA,^* and *Parp2^+/EA^* embryos. (**D**) The absolute number of I-IV cells in FLs from E13.5 *Parp2^+/+^, Parp2^-/-,^* and *Parp2^EA/EA^*embryos. (**E**) Representative IHC images of the FL from E13.5 *Parp2^+/+^*, *Parp2^-/^*^-^, and *Parp2^EA/EA^* embryos stained for TER119 and TUNNEL. The means ± SEM were shown for all the bar graphs, and an unpaired two-tailed t-test was used to measure the p values. ns, p>0.05; * p<0.05; **, p<0.01; ***p<0.001.

**Figure S3.**
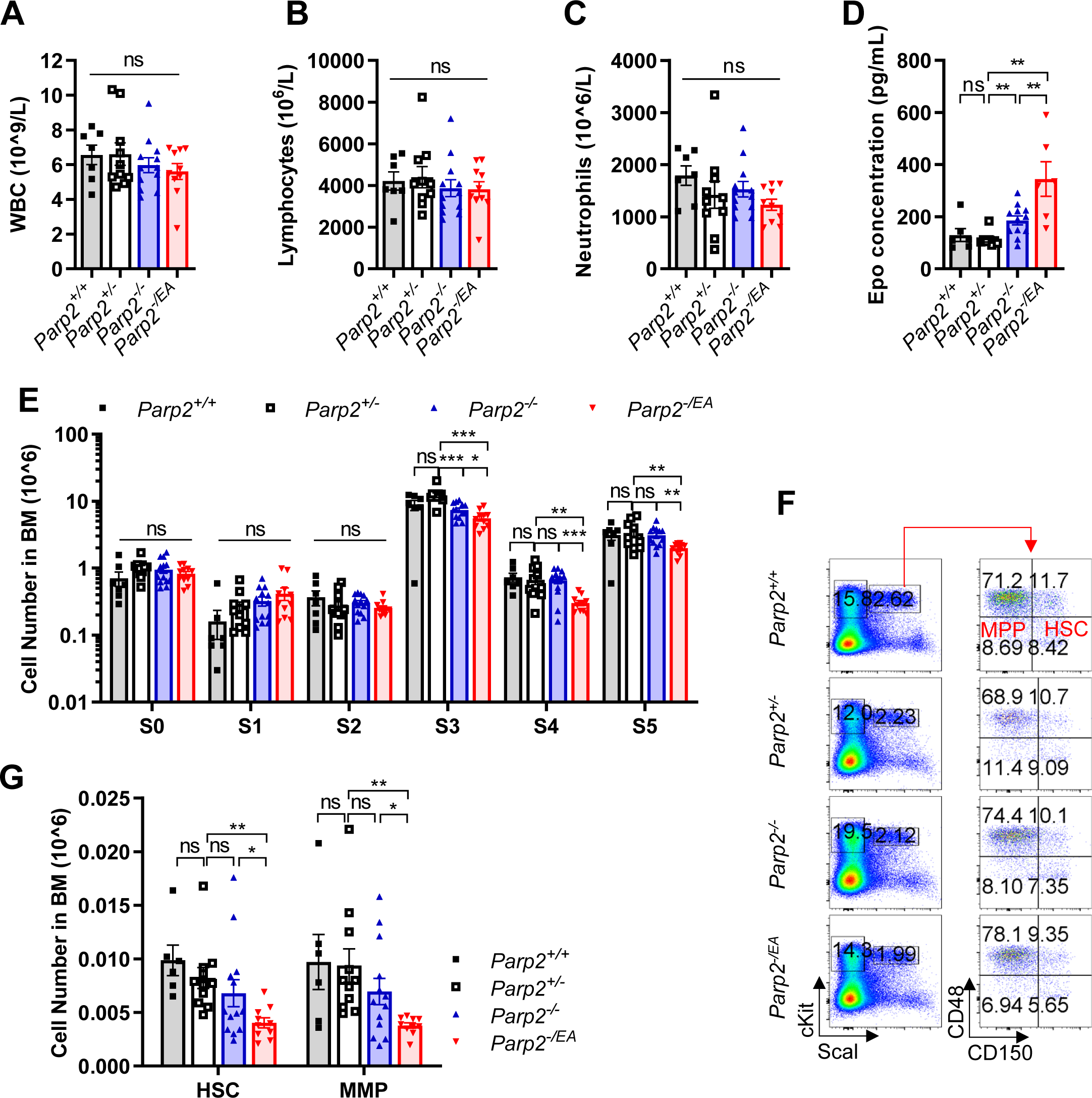
Hematopoiesis analysis of *Parp2^-/EA^* adult mice. (**A**) White blood cell count, (**B**) lymphocyte count, (**C**) neutrophil count and (**D**) Epo concentration in peripheral blood from *Parp2^+/+^, Parp2^+/-,^ Parp2^-/-,^*and *Parp2^-/EA^* mice. (**E**) The absolute number of S0-S5 cells in BM from *Parp2^+/+^, Parp2^+/-,^ Parp2^-/-^*and *Parp2^-/EA^* mice. (**F**) Representative flow cytometry analyses and (**G**) the absolute number of hematopoietic stem cells (HSC, Lin^-^Sca1^+^cKit^+^CD48^-^CD150^+^) and multi-potent progenitors (MPP, Lin^-^Sca1^+^cKit^+^CD48^-^CD150^-^) in BM from *Parp2^+/+^, Parp2^+/-,^ Parp2^-/-^* and *Parp2^-/EA^* mice. The means ± SEM were shown for all the bar graphs, and an unpaired two-tailed t-test was used to measure the p values. ns, p>0.05; * p<0.05; **, p<0.01; ***p<0.001.

**Figure S4.**
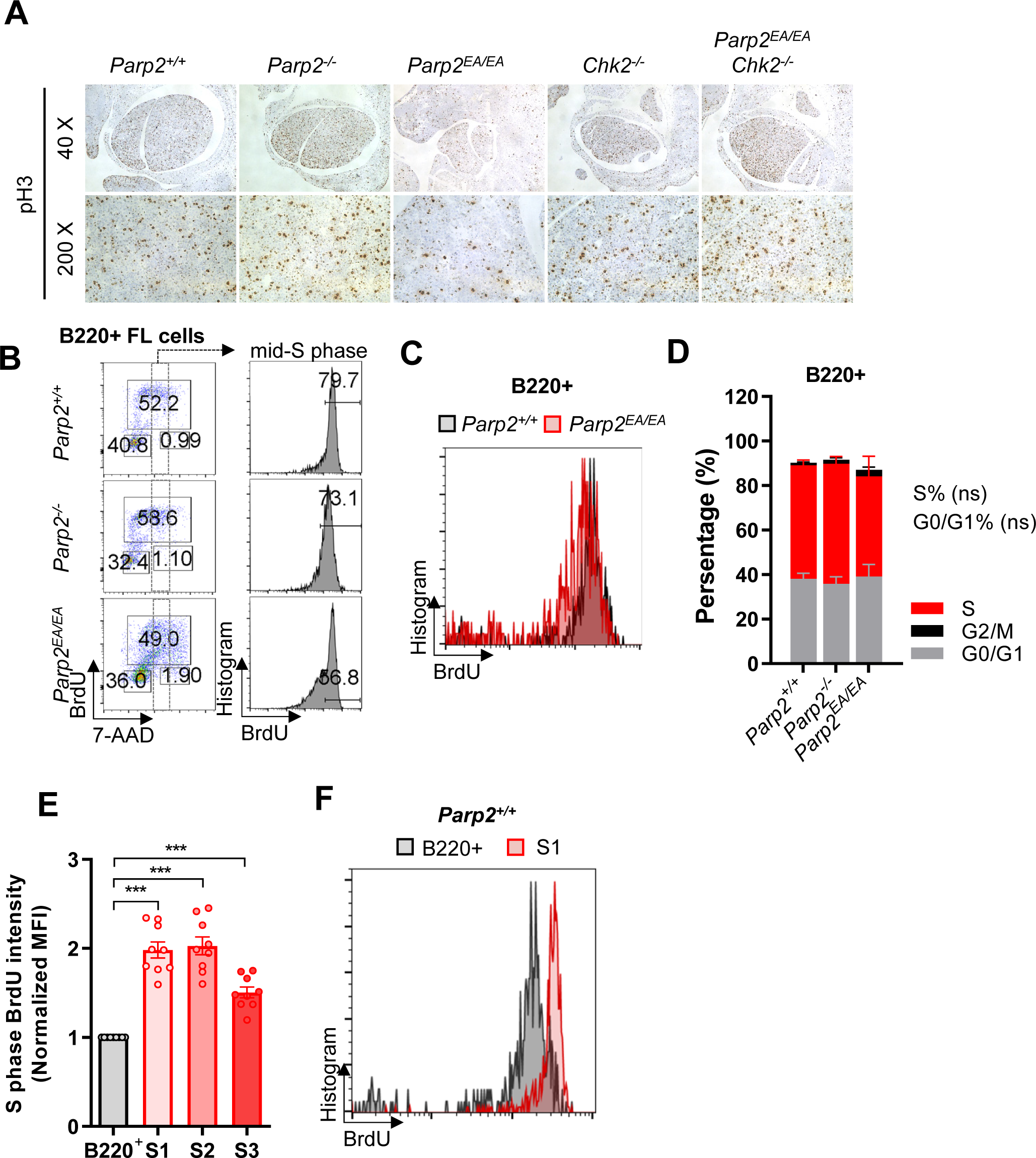
Phospho-Histone H3 staining and cell cycle distribution of B220^+^ cells in FL. **(A)** Representative IHC images of the FL from E13.5 *Parp2^+/+^*, *Parp2^-/^*^-^, *Parp2^EA/EA^*, *Chk2^-/-^* and *Parp2^EA/EA^;Chk2^-/-^* embryos stained for mitotic marker – Ser10-phospho-Histone H3 (pH3). **(B)** Representative flow cytometry analysis and (**D**) quantification of the cell cycle of B220^+^ cells from the FL of E13.5 *Parp2^+/+^, Parp2^-/-^* and *Parp2^EA/EA^*embryos. (C) Overlapping of BrdU levels in S phase B220+ FL cells from representative *Parp2^+/+^* and *Parp2^EA/EA^* embryos. **(E)** Normalized MFI of BrdU from B220+ FL cells and S1-S3 erythroblasts from FL. Each dot represents an individual embryo. (F) Representative flow cytometry analysis of the distribution of BrdU in mid-7AAD of B cells and S1 erythroblasts from the FL of E13.5 *Parp2^+/+^* embryos.

**Figure S5.**
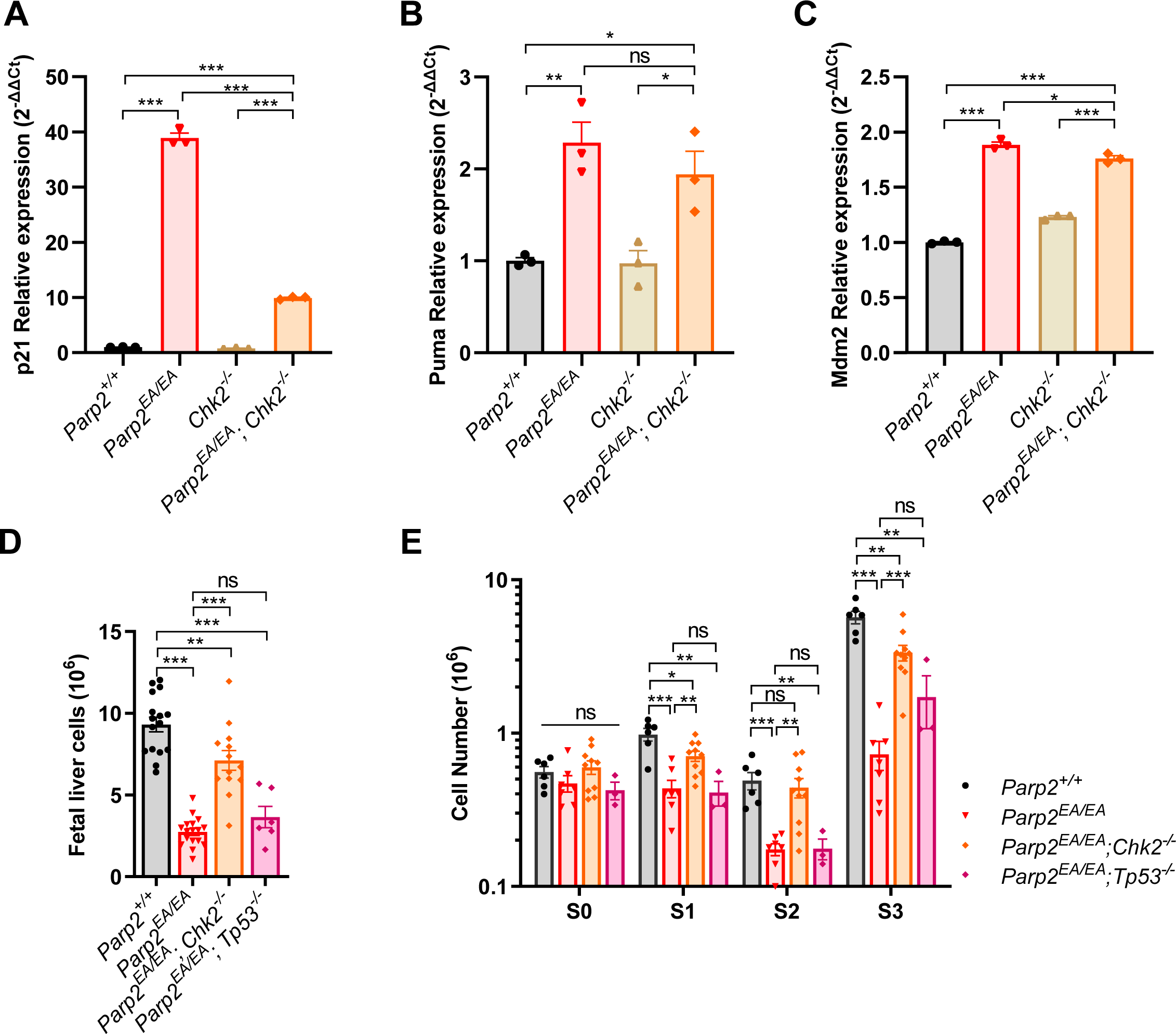
*Chk2* or *Tp53* deficiency rescues the embryonic lethality of *Parp2^EA/EA^* mice. (**A-C**) RT-qPCR to detect the relative expression of Tp53 target genes: (**A**) p21, (**B**) Puma and **(C)** Mdm2 in the FL cells of E13.5 *Parp2^+/+^*, *Parp2^EA/EA^*, *Chk2^-/-^* and *Parp2^EA/EA^; Chk2^-/-^* embryos. (**D**) Absolute number of FL cells from E13.5 *Parp2^+/+^, Parp2^EA/EA^*, *Parp2^EA/EA^; Chk2^-/-^* and *Parp2^EA/EA^;Tp53^-/-^* embryos. (**E**) Absolute number of S0-S3 cells in the FL from E13.5 *Parp2^+/+^, Parp2^EA/EA^*, *Parp2^EA/EA^; Chk2^-/-^* and *Parp2^EA/EA^;Tp53^-/-^* embryos. The means ± SEM were shown for all the bar graphs, and the unpaired two-tailed student’s t-test was used to measure the p values. ns, p>0.05; * p<0.05; **, p<0.01; ***p<0.001.

**Figure S6.**
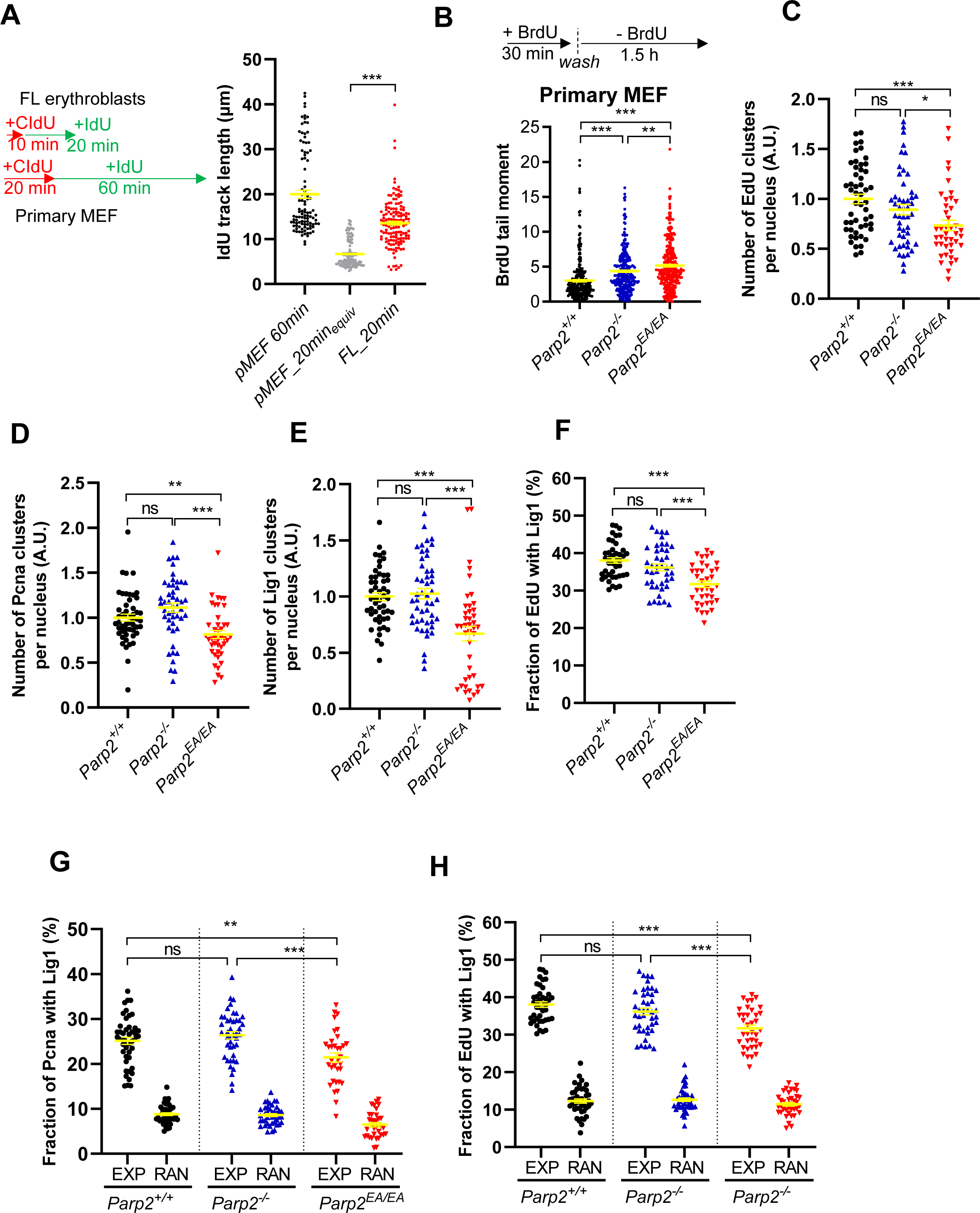
Quantification results of SMLM data. (**A**) A comparison of replication fork speed between *in vitro* cultured primary MEFs and FL cells. The IdU track length of *Parp2^+/+^* primary MEFs from Fig. 5C and *Parp2^+/EA^* FL cells from Fig. 4C were replotted side by side, with the 20 min equivlative IdU track length of primary MEFs plotted in grey dots. (**B**) BrdU comet tail moment in nascent DNA strands of primary MEFs from E13.5 *Parp2^+/+^, Parp2^-/-^* and *Parp2^EA/EA^*embryos. MEFs were pulse-labeled with 100 µM BrdU for 30 min and subsequent 1.5 h chase. BrdU comet tail moments were scored with anti-BrdU staining to label the nascent DNA strands. (**C-E**) Quantification of SMLM data for the relative number of (**C**) EdU, (**D**) Pcna, and (**E**) Lig1 clusters per nucleus of *Parp2^+/+^, Parp2^-/-,^* and *Parp2^EA/EA^* primary MEFs. Data were normalized to that of *Parp2^+/+^*primary MEFs. Individual data points represent values from a single nucleus. n > 40 nuclei were analyzed in three independent biological replicates. (**F**) Fraction of EdU associated with Lig1 within a 40 nm radius in *Parp2^+/+^, Parp2^-/-,^* and *Parp2^EA/EA^*primary MEFs. (**G**) The fraction of Pcna associated with Lig1 and (**H**) Fraction of EdU associated with Lig1 within a 40 nm radius were plotted alongside corresponding values obtained after cluster randomization (RAN). Individual data points represent values from a single nucleus. n>30 nuclei were analyzed in three independent biological replicates. For all the scatter dot graphs, the means ± SEM were shown, and an unpaired two-tailed t-test was used to measure the p values. p>0.05; * p<0.05; **, p<0.01; ***p<0.001.

**Figure S7.**
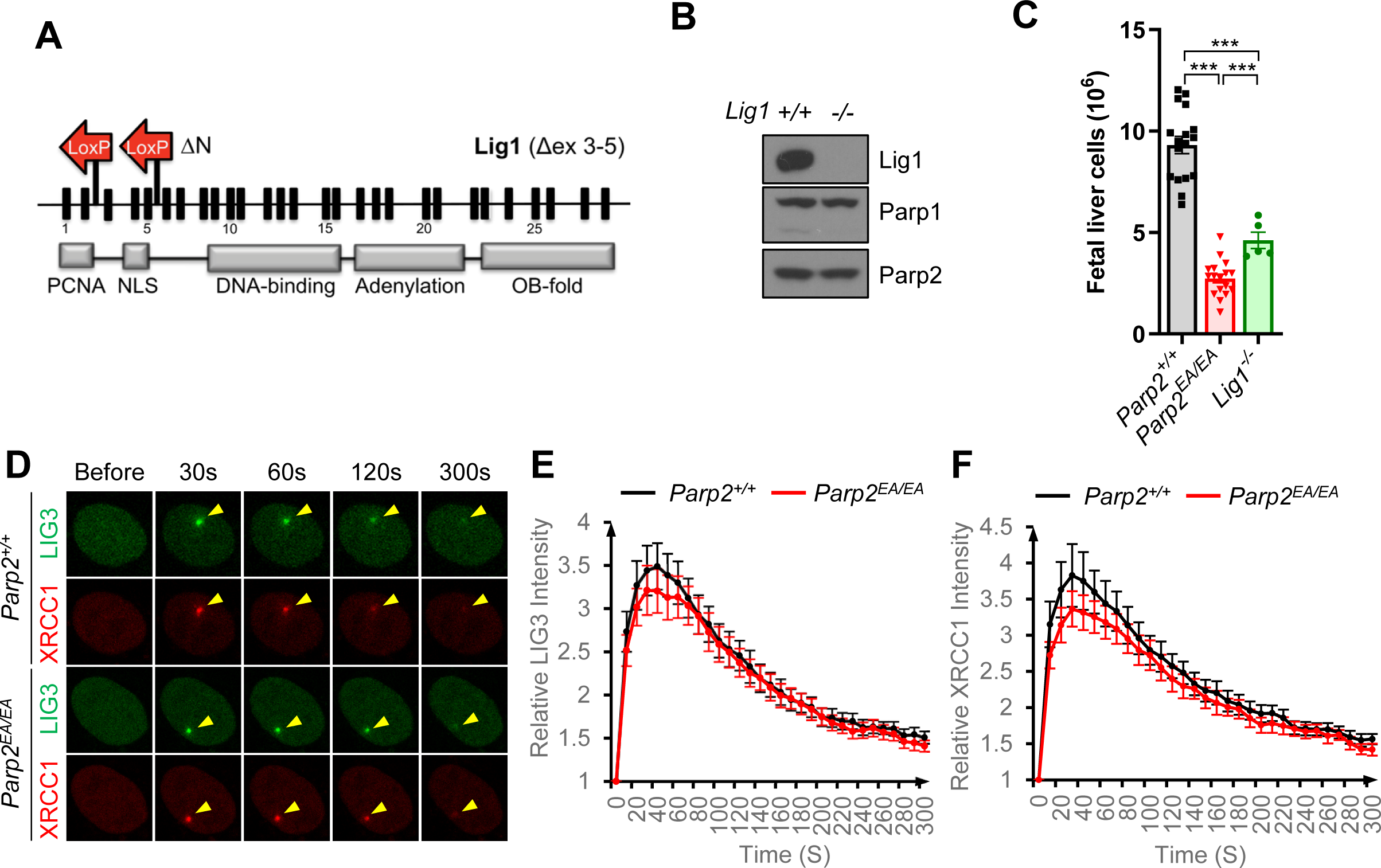
Generation of Lig1 deficiency mice. (**A**) Targeting scheme of Lig1-LoxP allele (Δex 3-5). The exons of Lig1 were shown as block boxes. The structural domains of Lig1 were shown as grey boxes. Two Red arrows show the location of LoxP on *Lig1*. (**B**) Western blot confirmed the deletion of Lig1 in iMEFs derived from E13.5 *Lig1*^-/-^ embryo. (**C**) Total cell count of FL from E13.5 *Parp2^+/+^, Parp2^EA/EA^* and *Lig1^-/-^* embryos. The means ± SEM were shown, and an unpaired two-tailed t-test was used to measure the p values. ***p<0.001. (**D**) Representative images of EGFP-LIG3 and mRFP-XRCC1 foci formation, and (**E and F**) the relative intensity kinetics of (**E**) EGFP-LIG3 and (**F**) mRFP-XRCC1 at DNA damage sites induced by 405 nm micro-irradiation. Data represent the means ± SEM from one representative experiment of three independent experiments with n > 7 cells each time with consistent results.

**Figure S8.**
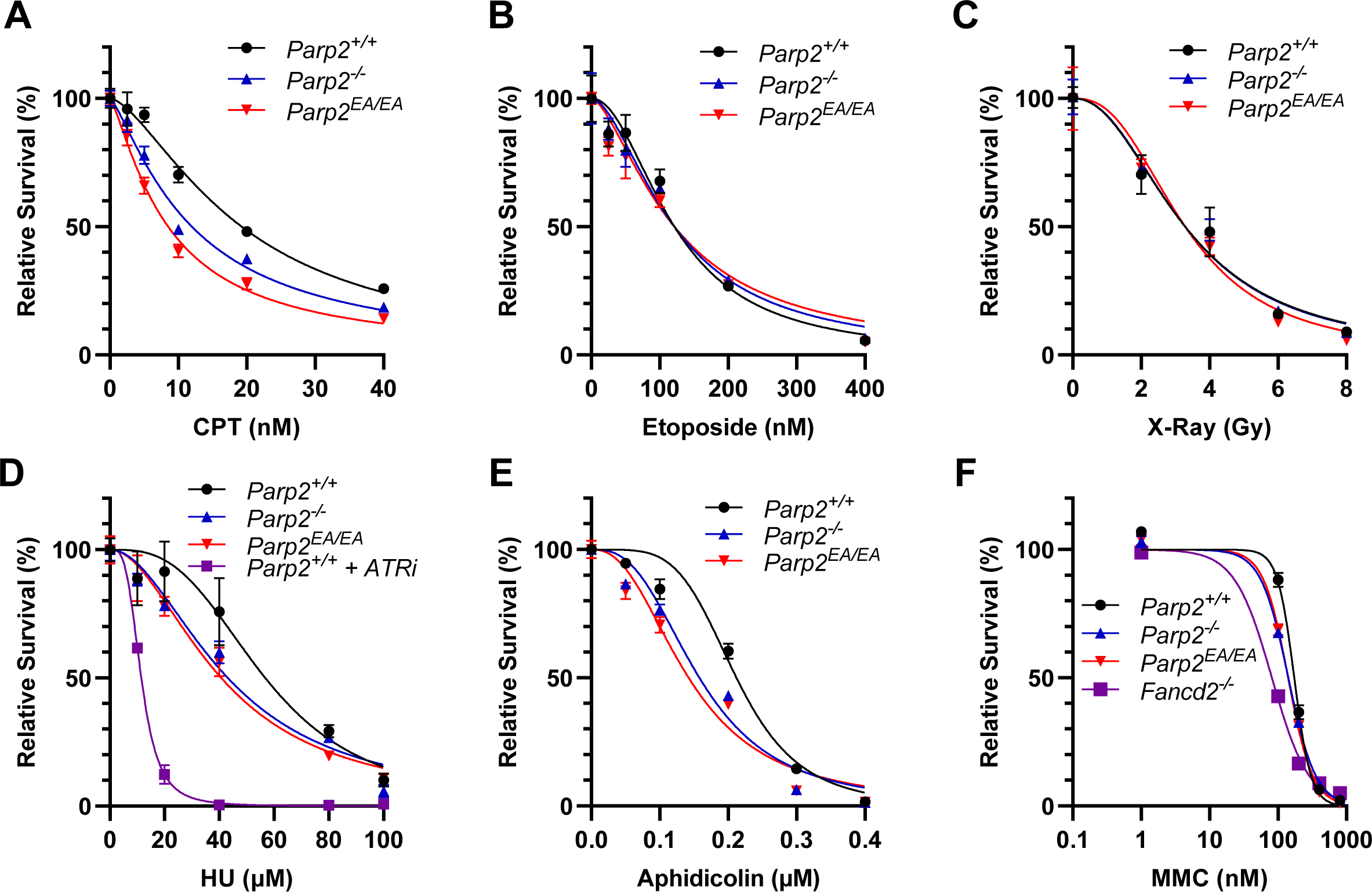
Inactive Parp2 sensitizes immortalized MEFs to various DNA damage. (**A-F**) Sensitivity assays of *Parp2^+/+^, Parp2^-/-^* and *Parp2^EA/EA^* iMEFs to (**A**) Camptothecin (CPT), (**B**) Etoposide, (**C**) X-Ray, (**D**) Hydroxyurea (HU), (**E**) Aphidicolin and (**F**) Mitomycin C (MMC). Cells were cultured in a medium containing indicated concentrations of DNA damage drugs for 5 days before conducting the MTT assay. In the case of the X-Ray sensitivity assay, cells were treated with indicated doses of X-Ray, then cultured in fresh medium for 5 days before the MTT assay. Data represents the means ± SD from one representative experiment out of three independent experiments. Each experiment included triplicate samples, and consistent results were observed across all three experiments.

**Figure S9.**
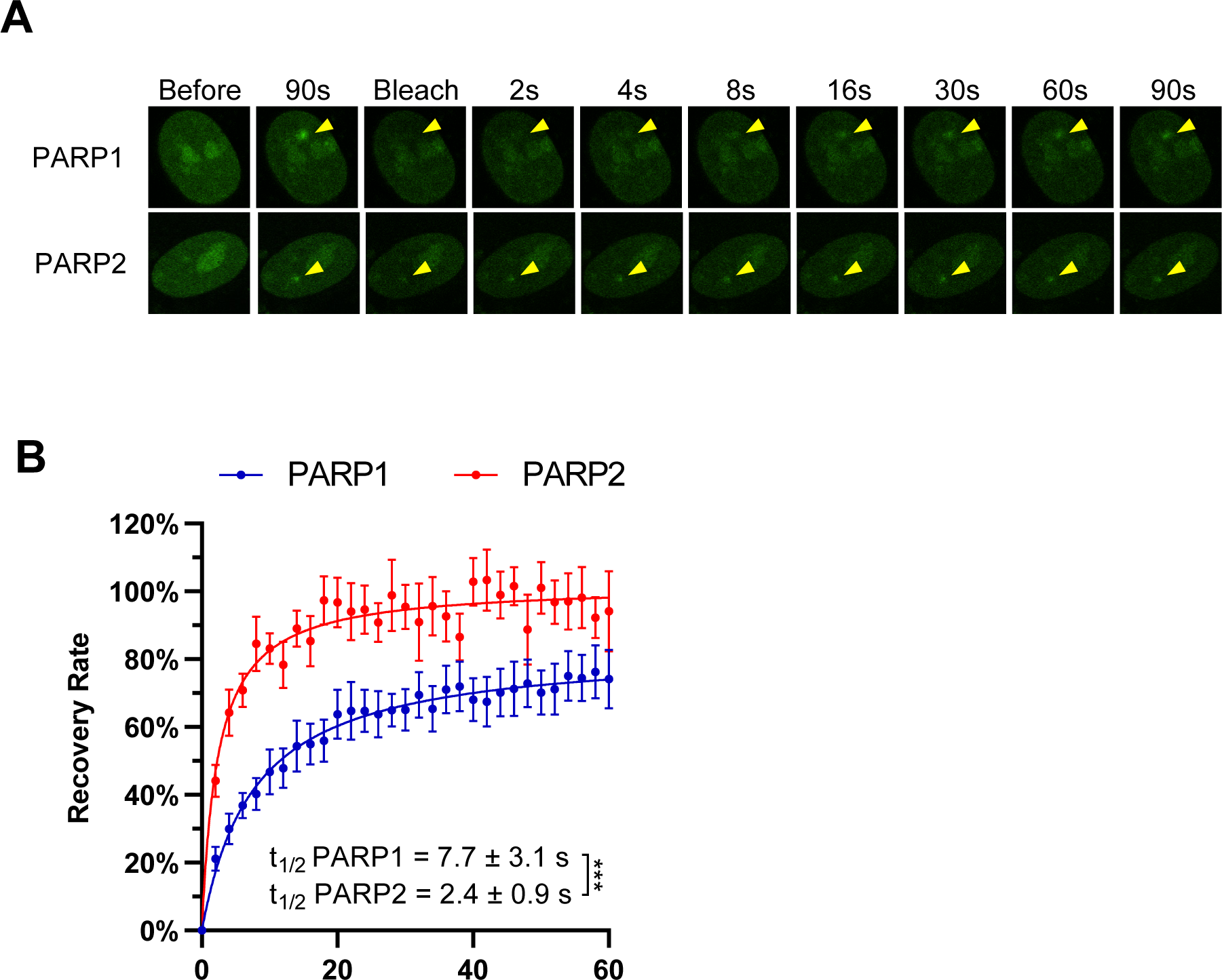
PARP1 and PARP2 FRAP assay in *PARP1/2 DKO* RPE-1 cells. (**A**) Representative images and (**B**) calculated FRAP recovery curve for EGFP-PARP1 and EGFP-PARP2 in *PARP1/2 DKO* RPE-1 cells. *t_1/2_* = 7.7 ± 3.1 s, *B_max_* = 83.5 ± 7.3% for PARP1; *t_1/2_* = 2.4 ± 0.9 s, *B_max_* = 102.0 ± 4.7% for PARP2. All the dots and bars represent means and SEM, respectively. P value was calculated using the extra sum-of-square F test. *** p<0.001.

